# Spatiotemporally dynamic noradrenergic regulation of cortical networks

**DOI:** 10.64898/2026.05.11.724286

**Authors:** Clayton Barnes, Felipe A. Cini, Peng Xu, Jessica A. Cardin

## Abstract

Brain activity and cognition exhibit state-dependent fluctuations that may reflect the influence of neuromodulatory systems, including norepinephrine (NE). Although the activity of noradrenergic neurons is strongly coupled to sleep-wake cycles and pupil dynamics, suggesting a global arousal signal, recent evidence suggests potentially modular partitioning of these cells. In addition, it is unclear whether different neuromodulatory signals exhibit distinct spatiotemporal patterns. We performed simultaneous, dual color mesoscopic imaging of either NE and calcium or NE and acetylcholine (ACh) signals across the neocortex, along with high-density electrophysiology, to investigate the relationship between NE release and neural activity. We find that the pattern of cortical NE signaling varies with behavioral state and is associated with enhanced functional connectivity. Simultaneous imaging of NE and ACh reveals spatiotemporally dynamic coupling between signals. Finally, NE signaling and functional connectivity are disrupted by sleep deprivation. Overall, our findings demonstrate that NE provides a multimodal signal that links transitions in behavioral state to cortical network interactions.

## Introduction

Animals cycle through multiple waking brain states ^1-4^ that are associated with different underlying brain network dynamics ^3, 5-11^ and regulated by neuromodulatory systems including norepinephrine and acetylcholine ^12, 13^. Both local circuits and large-scale networks spanning multiple cortical areas exhibit changes in coordination across behavioral state transitions. Norepinephrine (NE) released by neurons in the locus coeruleus (LC) plays a key role in non-REM sleep and sleep-wake transitions and is required to maintain wakeful patterns of cortical activity ^14, 15 16-20^. LC neurons exhibit changes in firing patterns with arousal and sensory stimulation ^15, 21, 22^ and the activity of LC axons tracks external markers of arousal, such as pupil dilation ^23, 24^. The noradrenergic system is thus thought to provide arousal-related signals throughout the cortex. However, the state-dependent dynamics of cortical noradrenergic release remain poorly understood. Moreover, the degree to which cortical noradrenergic signaling is coordinated with that of other key neuromodulatory systems, such as acetylcholine, is unknown.

Noradrenergic neurons in the LC receive input from and send projections to widespread cortical targets ^25^, suggesting that the LC-norepinephrine system may integrate broad inputs and provide a synchronized, homogeneous signal. However, evidence from targeted connectivity mapping has suggested LC neurons projecting to different areas may instead represent distinct functional modules ^26-29^. Individual LC neurons may project to preferred targets ^30^ and LC neurons projecting to different cortical targets may receive different amounts of synaptic input and exhibit different activity patterns ^28^. It thus remains unclear whether noradrenergic release across the cortex is homogeneous or spatiotemporally restricted.

Noradrenergic signaling regulates local circuit function via pre- and postsynaptic impacts on glutamatergic synapses^31^, shapes short-term synaptic plasticity^32^, and regulates GABAergic synaptic transmission ^33-35^. Ex vivo findings suggest that norepinephrine may depolarize both cortical excitatory neurons and GABAergic interneurons, increasing spontaneous activity and decreasing adaptation ^36-40^. In the intact cortex, exogenously applied norepinephrine has been reported to promote depolarization and spontaneous firing associated with arousal and locomotion ^41^, to decrease spontaneous spiking ^42^, or to exert a dose-dependent effect where modest levels of NE enhance and high levels suppress firing ^43-47^. Noradrenergic modulation of local circuit activity may likewise regulate sensory-evoked responses in a dose- and area-dependent manner ^42, 43, 48-51^. However, the detailed relationship between endogenous noradrenergic release and cortical activity remains unclear. Furthermore, the impact of norepinephrine on the coordinated activity of large-scale circuits is largely unknown.

The recent development of a suite of receptor-based fluorescent indicators that directly report neuromodulatory binding has made possible the exploration of neuromodulation across behavioral states and brain areas. Here, we used two-color mesoscopic imaging ^52-54^ across the entire dorsal neocortex of the awake mouse, in combination with high-density electrophysiology ^55, 56^, to quantify the relationships between behavioral state, cortical activity, and noradrenergic signaling. Our results demonstrate that NE release is state-dependent and exhibits multiple modes, including spatiotemporally heterogeneous release across cortical areas and brief, periodic epochs of homogeneous signaling. The spatiotemporal pattern of noradrenergic release is largely distinct from simultaneously recorded cholinergic dynamics. Cortical activity across scales is modulated with fluctuations in NE signaling, and pharmacological manipulation of noradrenergic receptor activity alters cortical functional connectivity. Finally, modest sleep deprivation leads to disruption of the normal dynamics of NE release during wakefulness and perturbation of cortical networks. Overall, these findings revise our view of NE from a global signal of arousal to a multimodal and spatially heterogeneous signal that links transitions in behavioral state to cortical network interactions.

## Results

### Dual-color mesoscopic NE and calcium imaging

To simultaneously monitor neuronal activity and noradrenergic signaling in the neocortex of awake mice, we expressed the red fluorescent calcium indicator jRCaMP1b^53^ and a green fluorescent norepinephrine sensor, GRAB_NE_ (Ne2m) ^57^ throughout the brain via neonatal injection of AAV vectors into the transverse sinus^10, 58^. This approach resulted in cortex-wide, uniform expression of both reporters (Figure S1), and we confirmed both *ex vivo* and *in vivo* that the GRAB_NE_ signal specifically reported noradrenergic signaling (Figure S1). We then performed mesoscopic imaging^52^ of both reporters through the intact skull of mice that were head-fixed and freely running on a wheel (Figure 1a, see Methods). Imaging was performed by strobing 575nm (jRCaMP1b), 470nm (GRAB_NE_), and 395nm (control) excitation light with an overall frame rate of 10 Hz per channel. GRAB_NE_ and jRCaMP1b images were co-registered, aligned to the Allen Common Coordinate Framework (CCFv3, Figure S1)^59^, and normalized (z-scored).

**Figure 1.**
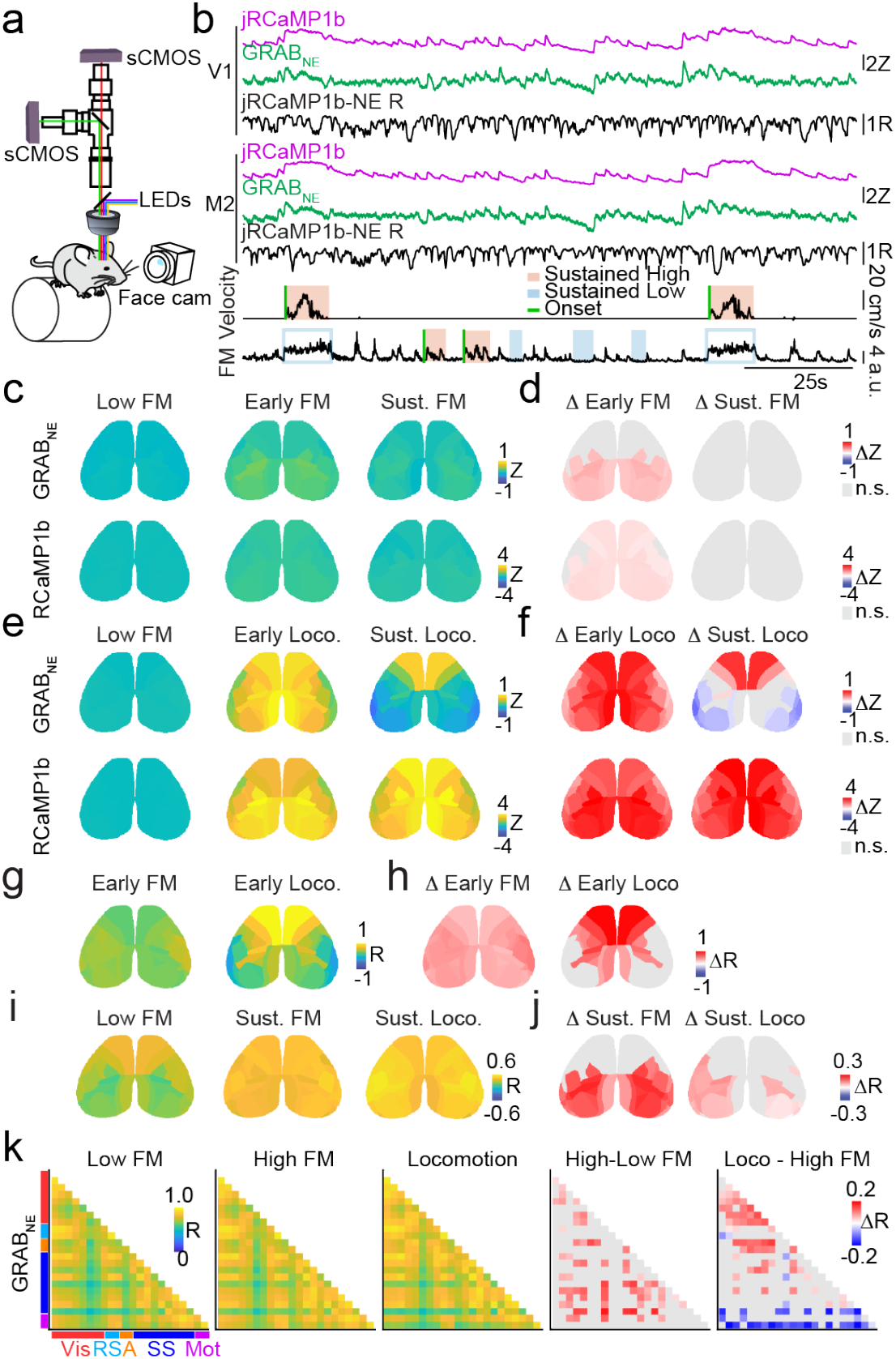
State-dependent spatiotemporal heterogeneity of cortical noradrenergic signals. **a**, Schematic of the dual-wavelength widefield imaging setup. **b**, Time series for GRAB_NE_ (green) and jRCaMP1b (magenta) signals in V1 and M2 parcels. Instantaneous (1.1 s window) Pearson’s correlation (R) between GRAB_NE_ and jRCaMP1b is shown beneath in black. Simultaneous facial motion and running speed are shown below. Example periods of sustained high (orange) and low (blue) states are highlighted for facial motion and locomotion. **c**, Average spatial maps (n=9 mice) showing z-scored GRAB_NE_ (upper) and jRCaMP1b (lower) activity at the onset of movement-defined behavioral states of facial motion (Early FM) and during sustained facial motion (Sust. FM). Trials were z-scored relative to a −5 to −2 s baseline. **d**, Change in signal during early and sustained facial motion compared to low facial motion for GRAB_NE_ (upper) and jRCaMP1b (lower). Stratified permutation testing (10K permutations) with Benjamini–Yekutieli FDR correction assessed significance of early and sustained facial motion epochs versus low facial motion; red indicates increases, blue indicates decreases, and gray indicates no significant change. **e**, Same as c, for locomotion. **f**, Same as d, for locomotion. **g**, Parcel-wise coupling between GRAB_NE_ and jRCaMP1b signals at the onset of facial motion (Early FM) or locomotion (Early Loco.). Coupling was quantified as the slope from a linear mixed-effects model. Raw slopes are shown as parcel maps. **h**, Change in signal coupling across states, gray indicates non-significant parcels. **i**, Parcel-wise correlations between GRAB_NE_ and jRCaMP1b signals during sustained low facial motion, high facial motion, and locomotion. **j**, Same as **h**, but for sustained states. **k**, Parcel-wise correlation matrices of GRAB_NE_ activity during sustained low facial motion, high facial motion, and locomotion states. Red indicates significantly increased correlations, blue indicates significantly decreased correlations, and gray indicates non-significance.

Both NE and calcium signals exhibited spatially heterogeneous, spontaneous fluctuations across the cortex (Figure 1b), illustrated by example data from two spatially distant cortical areas, primary visual (V1) and secondary motor (M2) cortex. Large fluorescence signal fluctuations occurred across cortical regions for both noradrenergic and calcium signals and co-varied with changes in behaviors such as locomotion and facial movement. We observed spatially varied activity patterns in both channels during periods of arousal and motor behavior (Figure 1b, highlighted time frames), suggesting that NE release is both dynamic across states and heterogeneous across cortical regions^57, 60^.

Periods of quiescence included epochs of both low and high facial movement, whereas locomotion always co-occurred with high facial movement, suggesting that arousal follows a progression from absence of motor activity, to facial motion, to locomotion ^3, 5^. We previously found that variation in brain activity across these distinct behavioral states is accompanied by both transient and sustained changes in cholinergic and calcium signals that reflect underlying circuit dynamics ^5^. Fluctuations in the NE signal were distinct during the onset of transitions to arousal states associated with facial motion and locomotion as compared to signaling during longer periods of these behaviors (Figure 1, Figure S1i-j). We therefore assessed NE and calcium signals during the early and sustained periods of facial motion and locomotion separately. Early, but not sustained, facial motion in the absence of locomotion was associated with a modest increase in both NE and calcium signaling (Figure 1c-d). In contrast, early locomotion was associated with spatially homogeneous NE and calcium signaling, whereas NE signaling during sustained locomotion periods was restricted to frontal cortical areas (Figure 1e-f). Noradrenergic signaling is thus state-dependent and exhibits distinct dynamics during state transitions compared to sustained periods of motor behavior. Statistical results for these and all subsequent analyses are listed in Supplemental Table 1.

Increases in noradrenergic signaling have been linked to regulation of local firing rates and modulation of neuronal sensitivity to inputs ^41, 43, 44, 46, 47^. We therefore investigated whether NE and calcium signals are coupled to each other within individual cortical areas and whether this relationship varies with behavioral state. Early facial motion was associated with a broad increase in correlation between the two signals (Figure 1g-h). In striking contrast, early locomotion was associated with a significant increase in inter-signal correlation that was largely restricted to the frontal cortex. In comparison, sustained periods of facial motion and locomotion were associated with enhanced correlations between the two signals that was spatially restricted to posterior cortical regions (Figure 1i-j). Overall, our findings demonstrate a spatially compartmentalized relationship between noradrenergic modulation and cortical activity that switches between modes with changes in behavioral state.

Behavior is thought to rely on the large-scale, coordinated activity of networks that span multiple cortical areas^61-63^. We therefore examined the spatiotemporal relationships of both NE and calcium signals by measuring pairwise correlations between cortical regions. We focused on sustained periods of low facial movement, high facial movement, and locomotion, allowing us to test whether these behavioral states with different average increases in fluorescence signals also correspond to distinct changes in network organization. We found that increased facial motion was associated with significant increases in between-area pairwise correlations across the cortex for the NE signal (Figure 1k). In accordance with our previous findings, facial motion was likewise associated with a general increase in pairwise cortico-cortical correlations in the calcium signal (Figure S1l). In contrast, locomotion was associated with an increase in correlations in the NE signal across posterior visual areas, but a reduction in the correlations between frontal and motor areas and the rest of the cortex (Figure 1k). Overall, our results suggest that moderate levels of arousal (i.e., facial movement without locomotion) generally enhance the coordination of NE release in the cortex, but further increases in arousal (i.e., locomotion) lead to spatially separated enhancement of NE signals in posterior areas and decorrelation of frontal NE signals.

### Modulation of cortical network activity

Norepinephrine is thought to regulate the firing of cortical excitatory ^36, 37, 40^ and inhibitory ^38, 64^ neurons. The activity of cortical neurons is enhanced at transitions between behavioral states (Figure S2e-f) ^3, 5^ that align with transient increases in neuromodulatory signaling (Figure 1, S1), suggesting that fluctuations in NE could be associated with increased firing of local neurons. To examine the organization of cortical activity around fluctuations in NE signaling, we performed simultaneous dual color mesoscopic imaging of GRAB_NE_ and RCaMP1b with high-density electrophysiology recordings using Neuropixels 2.0 probes ^56^ in V1 (Figure 2a, S2a-d). We found that the NE signal exhibited transient fluctuations in amplitude that were detectable as local peaks (Figure 2a), simultaneously observed across hemispheres, and associated with a homogeneous, pan-cortical release of NE (Figure 2c,d, S2j). Individual regular spiking (RS), putative excitatory neurons and fast spiking (FS), putative inhibitory interneurons exhibited transient increases in firing rate aligned to NE peaks (Figure 2b, S2 g-h). Overall, FS cells were more robustly modulated than RS cells (Figure 2c), and firing modulation decreased during locomotion as compared to lower arousal states (Figure S2f-g). The calcium signal, a measure of population activity, was enhanced around NE peaks across areas (Figure 2f, S2i) and this enhancement was larger and more spatially homogeneous with arousal and locomotion (Figure 2f). Together, these results suggest that cortical firing across layers and areas is modulated by transient global fluctuations in NE release.

**Figure 2.**
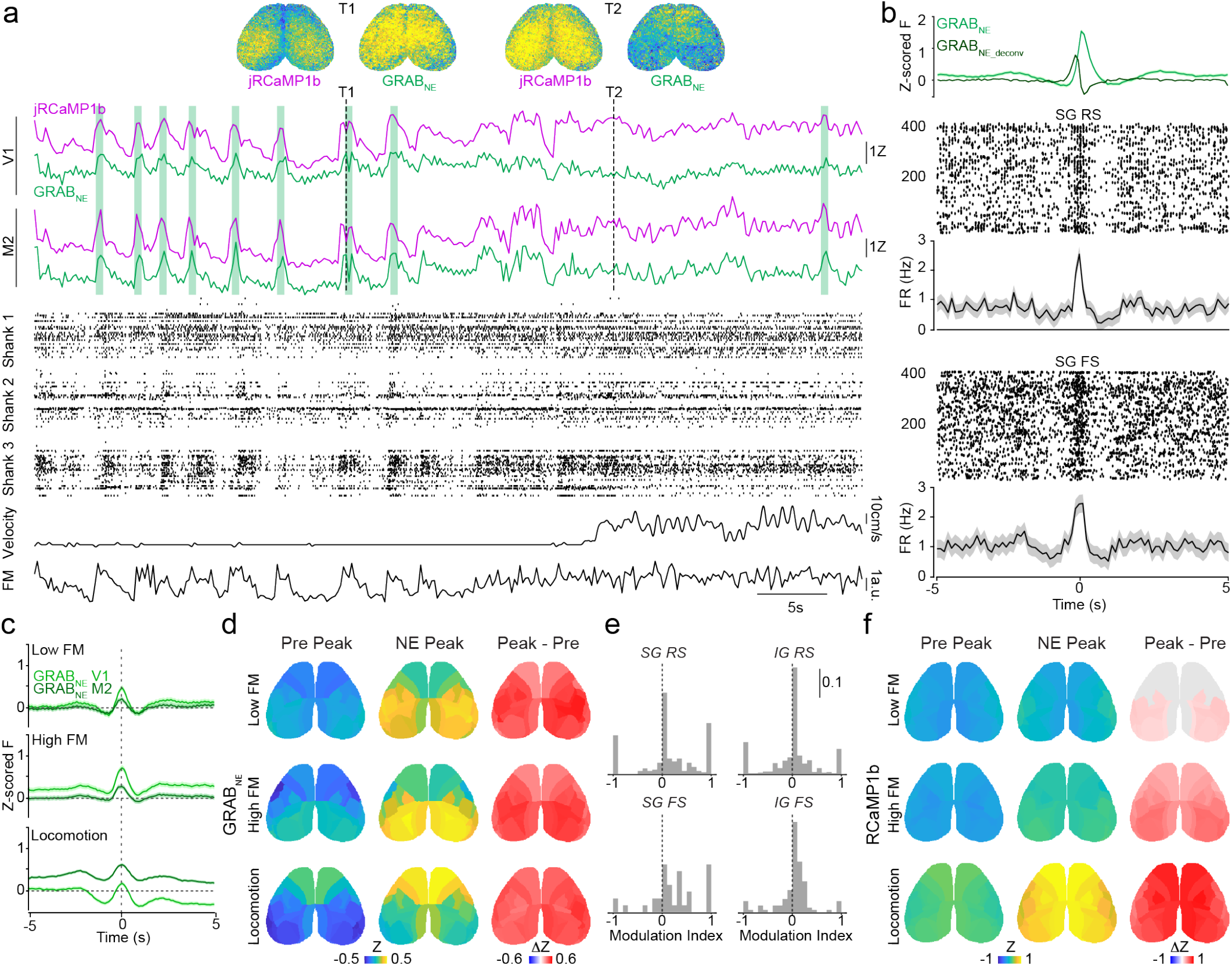
Fluctuations in norepinephrine release modulate cortical activity. **a**, Example GRAB_NE_ (green) and jRCaMP1b (magenta) traces aligned to single units recorded along a Neuropixels 2.0 probe in V1. Raster plots show spike times of units spanning three probe shanks, illustrating simultaneous mesoscopic imaging and high-density electrophysiology. Green shaded bars indicate detected local peaks in the GRAB_NE_ signal. Lower traces show facial motion and wheel velocity. Vertical dotted lines marking times T1 and T2 correspond to the snapshots above of activity in each channel at those times. **b**, Example single-unit raster plots around detected peaks in the local GRAB_NE_ signal (green) and GRAB_NE_ signal deconvolved using reported kinetics of the sensor (GRAB_NE_deconv_; dark green). Upper: Spike times for a supragranular (SG) regular-spiking (RS) unit. Lower: Spike times for a supragranular fast-spiking (FS) unit. **c**, Population average local GRAB_NE_ peaks in V1 (light green) and M2 (dark green), during periods of low facial motion (Low FM), high facial motion (High FM) and locomotion (n = 6 animals). **d**, Average GRAB_NE_ signal before (Pre Peak) and during (NE Peak) GRAB_NE_ peaks detected in V1. Corresponding significance maps (Peak-Pre) indicate parcels with significant differences between the two epochs, gray indicates non-significant parcels. Stratified permutation testing (10K permutations) with Benjamini-Yekutieli FDR correction (n = 6 animals). **e**, Modulation of the firing rates of supragranular (SG) and infragranular (IG) RS and FS cells around GRAB_NE_ peaks, shown as histograms of the modulation index values for each type (n = 6 animals, supragranular RS = 200, infragranular RS = 482, supragranular FS = 40, infragranular FS = 124). **f**, Average jRCaMP1b signal before (Pre Peak) and during (NE Peak) GRAB_NE_ peaks. Corresponding significance maps (Peak-Pre) indicate parcels with significant differences between the two epochs, gray indicates non-significant parcels. Stratified permutation testing (10K permutations) with Benjamini-Yekutieli FDR correction (n=9 animals).

Cortical circuits are simultaneously influenced by multiple neuromodulatory signals, which may have distinct state-dependent dynamics. We therefore performed dual color mesoscopic imaging of norepinephrine and acetylcholine in mice expressing the green fluorescent NE indicator GRAB_NE_ and a red fluorescent ACh indicator GRAB_ACh_ (rACh0.5h) ^65, 66^ throughout the cortex. Both the NE and ACh signals showed fluctuations that changed with behavioral state (Figure 3a). Moreover, the within-area correlation between the two signals was highly dynamic. Indeed, the fluctuations in NE and ACh exhibited periods of both high correlations at the transitions between behavioral states and distributed, lower correlations during periods of sustained arousal and motor activity (Figure 3b-f, S3a-b), suggesting dynamic multiplexing of neuromodulatory signals across the cortex. Simultaneous imaging of the two neuromodulators revealed distinct patterns of spatial correlations across cortical areas during sustained epochs of different behavioral states (Figure S3c-d), with ACh becoming spatially decorrelated ^5^ but NE showing spatially selective patterns of enhanced correlation during locomotion. Local peaks in the NE signal were associated with a decrease in ACh signaling when peaks occurred during periods of quiescence in the absence of facial motion (Figure 3g-h). In contrast, the peak of the NE signal co-occurred with global peaks in the ACh signal during periods of facial motion and locomotion, suggesting transient moments of spatially homogeneous signaling by both neuromodulatory systems during arousal. Together, these results suggest a spatiotemporally dynamic relationship between norepinephrine and acetylcholine release across the dorsal cortex, combining transient cortex-wide coordination with more distributed, state-dependent patterns that may enable flexible neuromodulatory control of cortical networks.

**Figure 3.**
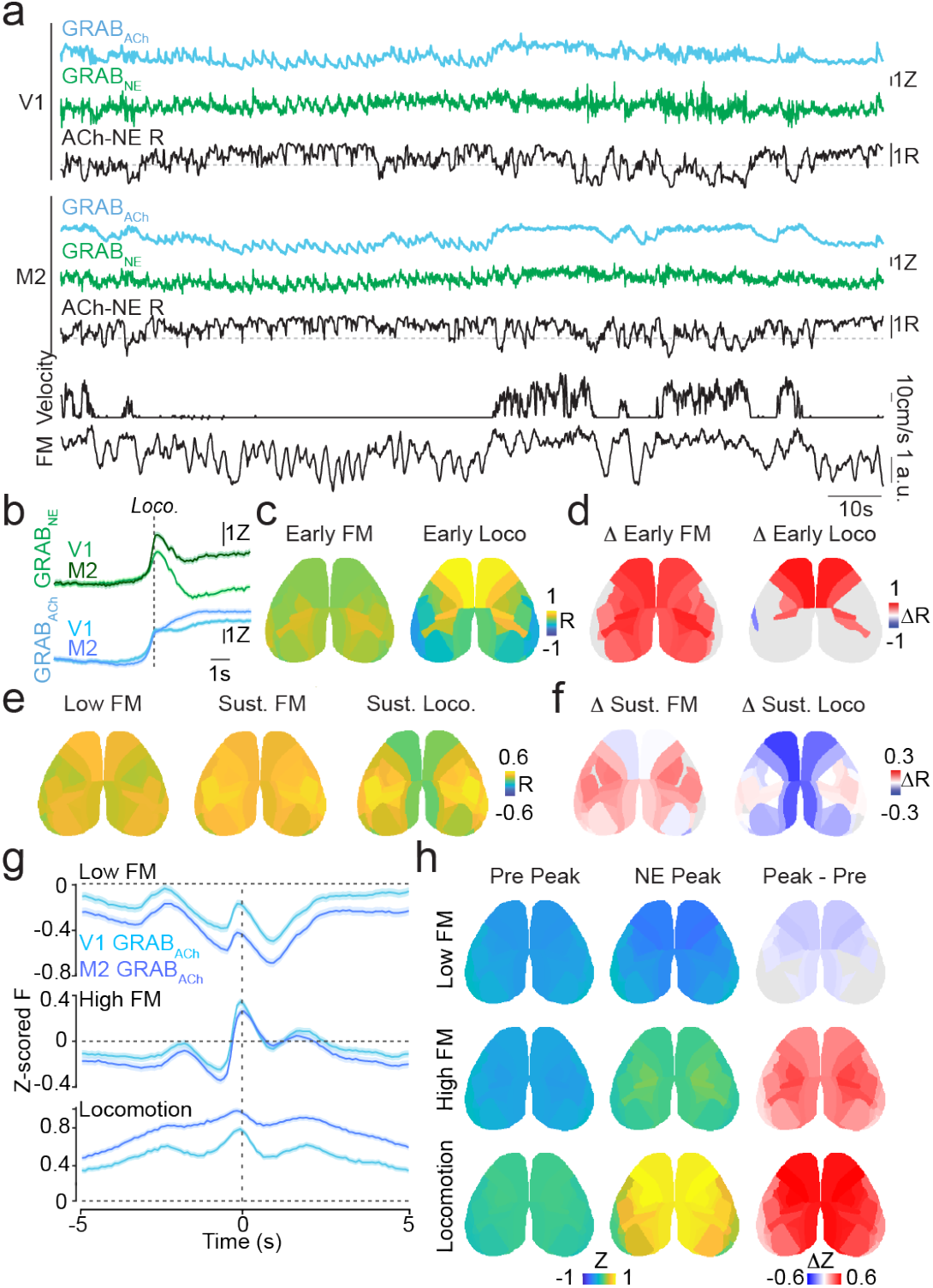
Region- and state-specific coupling of noradrenergic and cholinergic release. **a**, Example GRAB_NE_ (green) and GRAB_ACh_ (cyan) traces from V1 and M2 aligned to facial motion and running speed traces (black). Instantaneous correlations between the two signals are shown beneath each trace using a 1.1 s moving window (black). **b**, Average GRAB_NE_ and GRAB_ACh_ traces from V1 and M2 aligned to locomotion onset (dotted line). **c**, Parcel-wise correlations between GRAB_NE_ and GRAB_ACh_ signals at the onset of facial motion (Early FM) and locomotion (Early Loco.). **d**, Stepwise comparisons of sustained-state correlations, quantifying the results in **c**. Red indicates significantly increased correlations, blue indicates significantly decreased correlations, and gray indicates non-significance (n=6 animals, stratified permutation test, 10K permutations, Benjamini-Yekutieli FDR correction). **e**, Parcel-wise correlations between GRAB_NE_ and GRAB_ACh_ signals during sustained epochs of low facial motion (Low FM), high facial motion (High FM) and locomotion (Locomotion). **f**, Stepwise comparisons of sustained-state correlations, quantifying the results in **e**. Red indicates significantly increased correlations, blue indicates significantly decreased correlations, and gray indicates non-significance (n=6 animals, stratified permutation test, 10K permutations, Benjamini-Yekutieli FDR correction). **g**, Average GRAB_ACh_ traces in V1 (light blue) and M2 (dark blue) aligned to detected GRAB_NE_ peaks. **h**, Average GRAB_ACh_ signal prior to (Pre) and during (Peak) GRAB_NE_ peaks for low facial motion, high facial motion, and locomotion states and the change in signal (Peak – Pre). Red indicates significantly increased signal, blue indicates significantly decreased signal, and gray indicates non-significance (n=6 animals, stratified permutation testing (10k permutations) with Benjamini-Yekutieli FDR correction).

### Noradrenergic regulation of cortical functional connectivity

Like acetylcholine ^5^, norepinephrine may regulate both the pattern of local activity and cortico-cortical functional connectivity ^67-69^. To examine the impact of endogenous noradrenergic signaling on functional connectivity in cortical networks, we simultaneously recorded local and global cortical activity via a Neuropixels 2.0 probe and mesoscale RCaMP1b imaging, respectively, while pharmacologically blocking noradrenergic signaling in V1 (Figure 4a). Local blockade of α and β noradrenergic receptors led to overall decreased firing rates in deep layer RS cells, but not superficial RS or FS cells, compared to a vehicle control during sustained low and moderate arousal states (Figure 4b) but did not affect the cellular populations recorded (Figure S4a) or the behavioral state of the animal (Figure S4b). Spontaneous firing immediately preceding locomotion was diminished by blocking NE receptors, but the overall modulation of firing rates of individual RS and FS cells at state transitions was largely unaffected (Figure S4c-e)^41^. Noradrenergic receptor blockade in V1 significantly decreased the functional connectivity of V1 with other cortical areas, with long-range connectivity decreasing during both locomotion onset (Figure 4c) and sustained periods of each behavioral state (Figure 4d). Noradrenergic signaling thus regulates state-dependent activity in local cortical circuits and long-range functional interactions across cortico-cortical networks.

**Figure 4.**
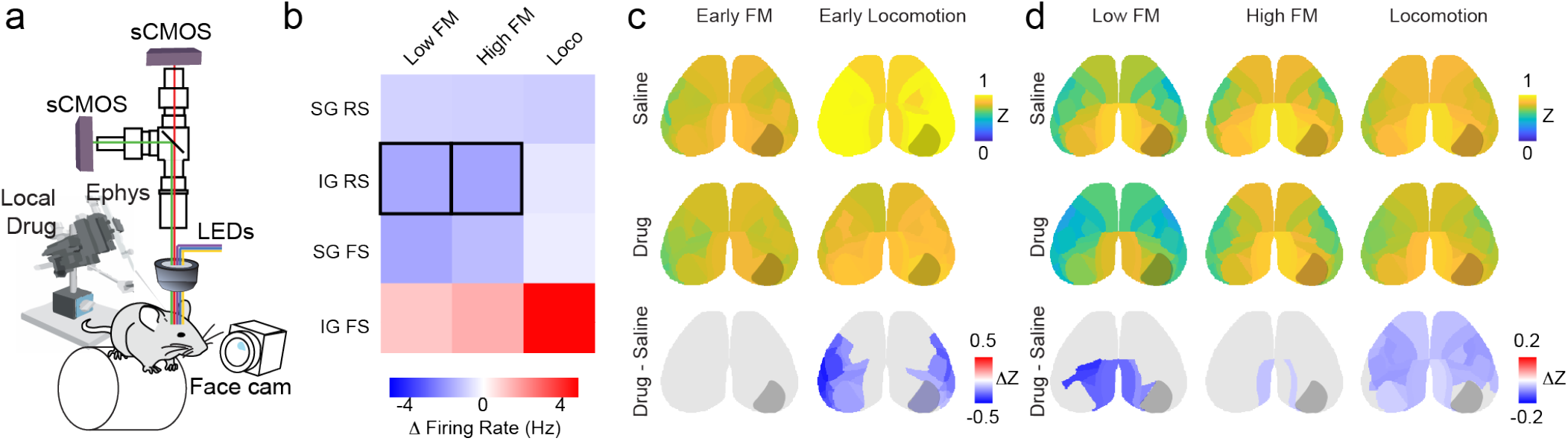
Noradrenergic blockade reshapes cortico-cortical functional connectivity. **a**, Schematic of mesoscopic imaging during local pharmacological manipulation. An adrenergic receptor antagonist cocktail (0.5 mM yohimbine, prazosin, and propranolol) was applied to V1 during simultaneous electrophysiological recording in V1 and mesoscale imaging of jRCaMP1b signals across the dorsal cortex. **b**, Change in firing rates for four cell classes (supragranular RS, supragranular FS, infragranular RS, infragranular FS) in each behavioral state (Low Facial Motion (Low FM), High Facial Motion (High FM), Locomotion (Loco)) upon application of the receptor antagonist cocktail, shown as a firing-rate modulation matrix. Red indicates increased firing rate, blue decreased firing rate, and black outlines indicate significant changes. (n = 6 mice, 500 IG RS, 235 SG RS, 123 IG FS, 47 SG FS cells, stratified permutation testing (10K permutations) with Benjamin-Yekutieli FDR correction). **c**, Correlation of drug-infused region of interest, right V1, to the rest of dorsal cortex during early facial motion and early locomotion. Lower plots (Drug – Saline) show difference in correlation between conditions (n=6 animals, stratified permutation testing (10K permutations) with Benjamin-Yekutieli FDR correction). Red indicates increased correlation, blue indicates decreased correlation, and gray indicates no significant change. **d**, Correlations between the right V1 and each cortical area during sustained periods of facial motion and locomotion. Lower plots (Drug – Saline) show difference in correlation between conditions (n=6 animals, stratified permutation testing (10K permutations) with Benjamin-Yekutieli FDR correction).

To examine whether norepinephrine similarly regulates the functional connectivity of individual neurons, we assessed the relationship between the spiking of RS and FS units in V1 and the RCaMP signal in other cortical areas. Noradrenergic receptor blockade decreased the functional connectivity of V1 neurons with both nearby and distant cortical regions (Figure 5a). We next examined the influence of noradrenergic signaling on the relationship between local cortical neurons and common spatiotemporal motifs of activity across cortical areas. Using SeqNMF ^70^, we identified four distinct, repeating spatiotemporal motifs of activity across the cortex that were observed in all animals (Figure S5a-c). Motif occurrence was not modulated by behavioral state (Figure S5d-f), suggesting that the motifs represent core patterns of activity arising from underlying cortico-cortical network architecture. Using a linear mixed-effects regression relating neuron–motif coupling effect sizes to noradrenergic modulation of individual neurons, we found that the degree to which a neurons’ firing rate was modulated by NE signaling was predictive of that cell’s association with specific cortical motifs (Figure 5b, S5g). Local blockade of noradrenergic receptors in V1 did not affect motif occurrence (Figure S5h), but significantly reshaped the relationship between individual RS and FS neurons and the four cortical activity motifs (Figure 5c). Acute manipulation of local noradrenergic signaling thus reorganizes the functional connectivity of individual V1 neurons and their relationship with common large-scale cortical activity patterns, further supporting a role for NE in long-range functional interactions.

**Figure 5.**
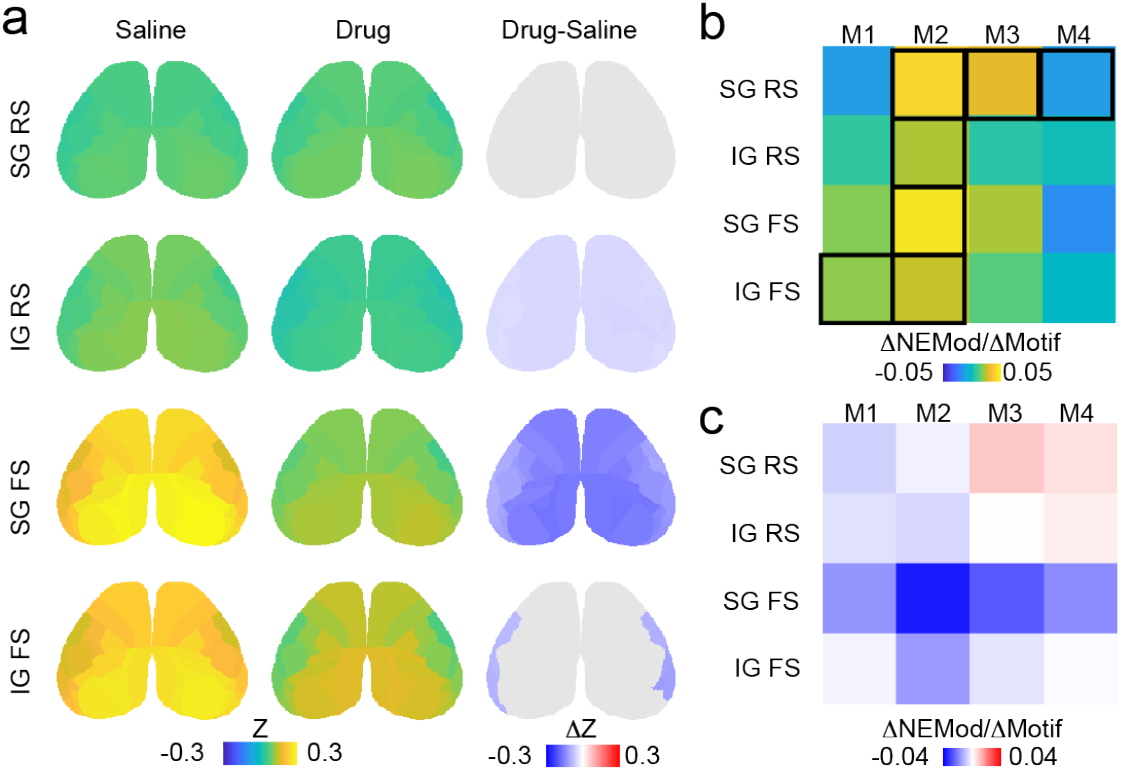
Noradrenergic modulation of interactions between single neurons and large-scale networks. **a**, Parcelwise correlations with the firing of individual units in V1 under Saline (left) and Drug (center) conditions. Change in unit-area correlations between conditions (Drug – Saline) is shown to the right. Red indicates significantly increased correlations, blue indicates significantly decreased correlations, and gray indicates non-significance (n=846 cells in 6 mice, stratified permutation test, 10K permutations, Benjamini-Yekutieli FDR correction). **b**, Strength of association between the noradrenergic modulation of the four cell classes (supragranular RS, supragranular FS, infragranular RS, infragranular FS) and each of the four identified cortical motifs (Motifs 1-4), measured as ratio of Cohen’s D (Δ_NEMod_/Δ_Motif_). Black outlines indicate significance. **c**, Change in the association between modulation of individual cells and cortical motifs when a noradrenergic antagonist cocktail was applied locally to V1. Red indicates increased association, blue indicates decreased association, and white indicates no change. (n = 846 cells in 6 mice; log-odds likelihood-ratio test, p=0.008).

Noradrenergic signaling is strongly coupled to the sleep-wake cycle ^68, 71-73^ and previous work has suggested that sleep deprivation may lead to sustained changes in the release of NE ^74-78^. We therefore examined the impact of sleep deprivation on cortical noradrenergic signaling and network activity. Mice were imaged on each day of a four-day paradigm in which they were prevented from sleeping during the light phase of the circadian cycle for two days (Figure 6a-b). Sleep deprivation resulted in selectively elevated amplitudes of NE signaling across much of the cortex during sustained locomotion (Figure 6c-f, S6a-h), suggesting that sleep loss leads to sustained elevation of noradrenergic release during arousal. The rate of local NE peaks was unaffected by sleep deprivation (Figure S6m) as was the entrainment of RCaMP signals to NE peaks (Figure S6n-o), suggesting that sleep deprivation did not disrupt transient fluctuations in NE release. However, noradrenergic signaling exhibited a state-dependent spatiotemporal reorganization following sleep deprivation, with NE signals becoming less coordinated in frontal areas during periods of quiescence but broadly more coordinated during locomotion (Figure 6g, S6i). Sleep deprivation had no impact on the amplitude of calcium signaling across behavioral states (Figure 6d,f). However, calcium signals exhibited enhanced long-range correlations between frontal and posterior cortical areas during quiescence and broadly decreased inter-areal correlations during locomotion following sleep deprivation (Figure 6h). One day of post-deprivation recovery sleep was largely sufficient to return cortical noradrenergic signaling and network activity to pre-deprivation states (Figure S6e-h, k-l). To examine the impact of sleep deprivation on the relationship between NE release and calcium signaling across the cortex, we computed the within-area correlation between the two signals in each behavioral state. Sleep deprivation disrupted the coupling of NE signals and cortical activity in a state-dependent manner, leading to enhanced correlations selectively in a subset of frontal and posterior areas during sustained periods of locomotion. Together, these results suggest that sleep deprivation leads to disruption of the normal spatiotemporal dynamics of NE release in the cortex and state-dependent perturbation of long-range cortico-cortical network interactions.

**Figure 6.**
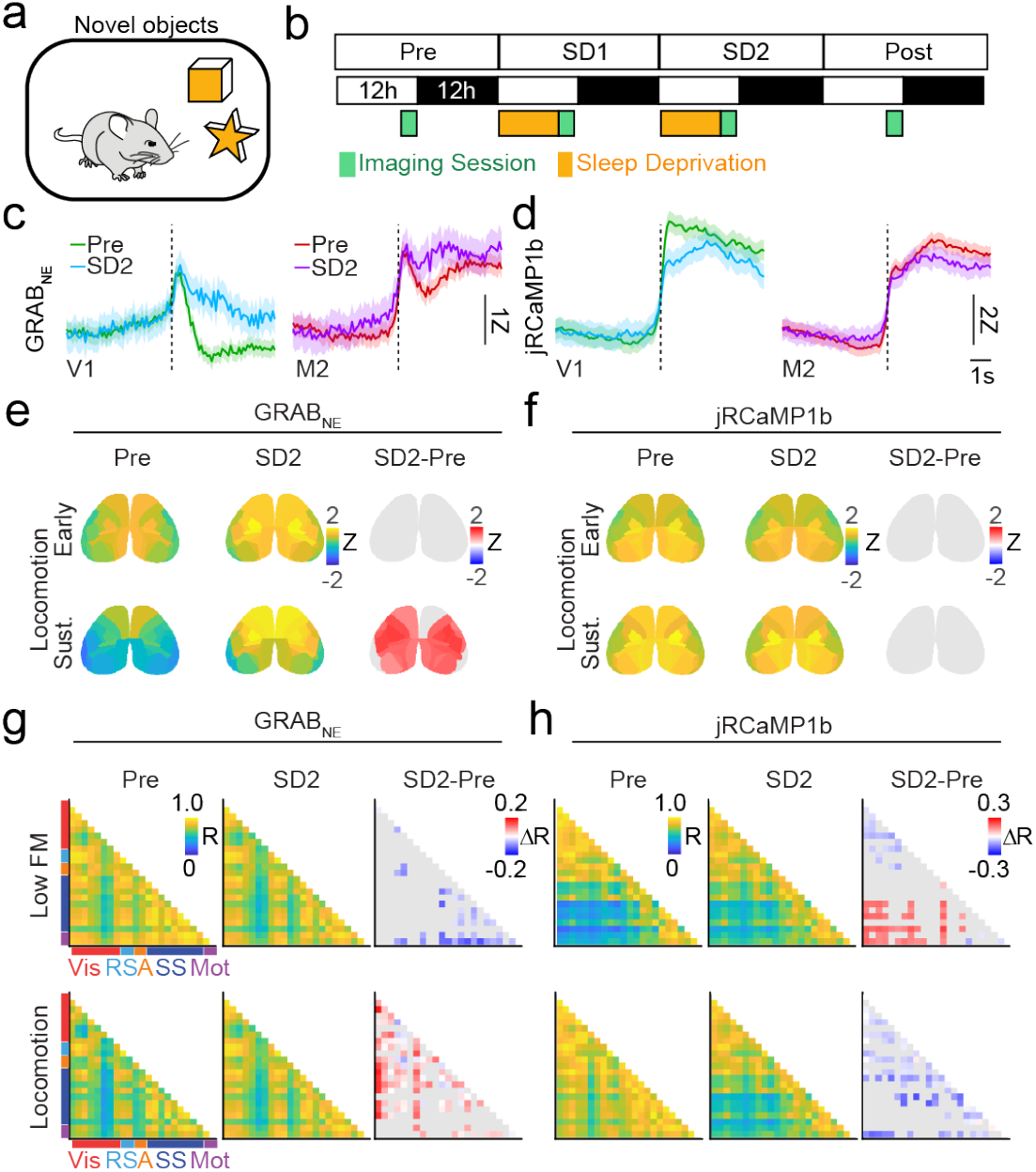
Sleep deprivation disrupts noradrenergic dynamics and cortical functional connectivity. **a**, Schematic of the novel object paradigm used to induce sleep deprivation. **b**, Experimental timeline, including a baseline imaging day (Pre), two consecutive sleep deprivation days (SD1, SD2), and a recovery day (Post). During sleep deprivation days, mice were prevented from sleeping during the light portion of the circadian cycle and allowed to sleep ad libitum during the dark portion of the cycle. **c**, Average GRAB_NE_ traces at locomotion onset (dotted line) from V1 (left) and M2 (right) on the second day of sleep deprivation (blue, purple) compared to baseline (green, magenta). **d**, Average jRCaMP1b traces at locomotion onset (dotted line) from V1 (left) and M2 (right) on the second day of sleep deprivation (blue, purple) compared to baseline (green, magenta). **e**, Dorsal cortex maps showing the amplitude of GRAB_NE_ signals during early (upper) and sustained (lower) periods of locomotion on days Pre and SD2, along with significance-thresholded maps of the change across days (SD2-Pre). (n=5 animals, stratified permutation testing (10K permutations) with Benjamin-Yekutieli FDR correction). **f**, Same as in e, but for jRCaMP1b signals. **g**, Pairwise correlations of GRAB_NE_ signals across cortical areas during sustained periods of low facial motion (Low FM) and locomotion on days Pre and SD2. Changes in the pairwise correlations are shown to the right (SD2-Pre). Increased correlations are denoted in red, decreased correlations are denoted in blue, and non-significant changes are denoted in gray. (n=5 animals, stratified permutation testing (10K permutations) with Benjamin-Yekutieli FDR correction). **h**, Same as in **g**, but for jRCaMP1b signals.

## Discussion

Locus coeruleus firing rates and calcium activity in the axons of LC projection neurons are strongly coupled to sleep-wake transitions and track pupil diameter, suggesting that the noradrenergic system may provide a global signal of arousal state ^14-24^. However, varied reports on the projection specificity of LC neurons suggest the potential for modularity in cortical noradrenergic signaling ^25-29^. Using a combination of mesoscopic imaging, high-density electrophysiology, and reporters for NE and ACh in awake behaving mice, we therefore examined the spatiotemporal dynamics of NE release across the dorsal cortex. Our results reveal that noradrenergic signaling in the cortex is multimodal, exhibiting spatiotemporal fluctuations that are related to distinct behavioral states. In addition, the coupling of NE to cortical network activity and the coordination between NE and ACh, another key neuromodulatory system, vary across cortical areas and behavioral states, suggesting a highly dynamic and area-specific relationship between state, neuromodulatory signaling, and cortical functional connectivity.

We found that noradrenergic signaling across the cortex was spatiotemporally diverse and varied across behavioral states associated with distinct motor behaviors. NE release was only modestly elevated during early and sustained periods of moderate arousal associated with facial motion. In contrast, noradrenergic signaling was spatially homogeneous during the onset of locomotion and transitioned to a spatially restricted area of release in frontal cortex during sustained periods of locomotion. In parallel, the spatial coordination of NE release across areas also varied across behavioral states, showing broadly enhanced inter-areal correlations during facial motion but reduced and enhanced correlations upon locomotion in distinct modules of frontal and posterior cortical regions, respectively. We further found that the noradrenergic signal exhibited intermittent fluctuations that were detected as spatially homogeneous peaks, potentially representing moments of global neuromodulatory influence over cortical networks. LC neurons exhibit periods of tonic and bursting activity that may differentially affect network function ^13, 21, 79, 80^, suggesting a possible model where distinct patterns of LC activity are linked to these modes of NE release and cortical network engagement.

Notably, the state-dependent dynamics of NE release were distinct from those we previously observed for the release of acetylcholine during the same behavioral states ^5^. Previous work has identified strong coherence between fluctuations in NE and ACh in prefrontal cortex that regulate cognitive control ^81^. However, other work has suggested that the activity of cholinergic and noradrenergic axons projecting to the cortex may exhibit distinct arousal-related dynamics ^24, 82-84^. We found that NE and ACh release was highly correlated at the onset of facial motion and locomotion, but less so during sustained periods within each behavioral state, suggesting that these two neuromodulatory systems may be briefly coupled at state transitions. Consistent with our previous findings on the spatial distribution of cholinergic signaling^5^, we observed coupling between noradrenergic and cholinergic signaling selectively in frontal cortical areas. Spatially broad peaks of noradrenergic signaling occurring during facial motion and locomotion, but not quiescence, coincided with spatially homogeneous cholinergic signaling. Our results thus suggest that noradrenergic and cholinergic signals exhibit specific, state-dependent moments of spatially selective coupling, as well as transient events of jointly homogeneous signaling across the dorsal cortex. Overall, these findings support the idea that multiplexed neuromodulatory signals may regulate cortical circuits ^85, 86^, providing functional flexibility to networks by influencing neural activity, synaptic release, and input integration under a wide range of modulatory regimes.

Previous work has suggested that noradrenergic signaling may have dose- and area-dependent effects on local neuronal activity ^42, 43, 48-51^. In particular, noradrenergic signaling may have distinct effects on frontal compared to sensory cortical circuits ^42, 43, 50, 51^. We found that NE signaling was differentially coupled to RCaMP signals in frontal and posterior areas across behavioral states, suggesting that the impact of NE release on local populations may vary across cortical areas. However, intermittent peaks of NE release across the cortex were associated with spatially broad activation of neural activity, indicating that transient high levels of NE signaling may uniformly increase overall population activity. Indeed, we found that the firing rates of many putative excitatory and inhibitory neurons across cortical layers in V1 were enhanced by transient fluctuations in endogenous NE signaling. In turn, local blockade of noradrenergic receptors decreased overall spontaneous firing rates, particularly at locomotion onset. Blockade of NE signaling did not affect modulation of firing by behavioral state transitions, indicating that NE may be only one of several signals regulating state-dependent modulation of cortical activity. Consistent with previous reports^41, 87^, both fast and regular spiking neurons were robustly modulated by noradrenergic signaling, suggesting that NE regulates the activity of both excitatory and inhibitory neurons in the local circuit. Together, our results suggest that the range of endogenous NE release observed across spontaneous behavioral states is associated with robust, area-specific modulation of neuronal activity across scales in the cortex. However, our paradigm did not include states of sleep, extreme arousal, or high stress, which may be associated with distinct regimes of NE release and neuronal modulation ^88^.

In addition to regulating the firing of neurons within local circuits, neuromodulatory signaling may modulate long-range functional connectivity between brain areas ^5, 67-69^. Indeed, we previously found that cholinergic signaling separately regulates local and long-range functional connectivity in a state-dependent manner^5^. Previous studies found that stimulation of the LC enhances functional connectivity between frontal and association areas ^89^ or globally across brain areas ^90^. Local NE signals in frontal cortex are also associated with variation in long-range functional connectivity based on fMRI signals^67^. We found that local blockade of noradrenergic receptors in V1 diminished functional connectivity between V1 and other cortical areas at both the population and single-cell levels. The dynamic spatiotemporal pattern of NE release may thus contribute to flexible modulation of long-range functional interactions across cortical areas.

Fluctuations in norepinephrine release are strongly coupled to sleep-wake cycles ^68, 71-73^, and low NE release is required for transitions between epochs of NREM and REM sleep ^17, 18, 71, 91^. Previous work in humans and animal models has suggested that sleep deprivation may tonically upregulate NE release ^74-78^. Using a paradigm for modest sleep deprivation, we found that loss of sleep led to enhanced levels of NE release and dysregulated the spatiotemporal structure of NE release across the cortex. In association, we observed a loss of the canonical structure of state-dependent functional connectivity in large-scale cortical circuits ^92-96^, suggesting that short-term sleep deprivation may result in a disrupted pattern of NE release and a substantial change in the neuromodulatory regulation of state-dependent network coordination. Long-term sleep loss, a common consequence of shift work ^97-99^, may potentially exacerbate these effects but remains to be studied further.

Our findings provide novel evidence that noradrenergic signaling in the cortex is spatially structured and plays a dynamic role in regulating circuit interactions across scales. Noradrenergic regulation of flexible circuit function during wakefulness varies with behavioral state, exhibits multiple spatiotemporal modes, and is highly sensitive to perturbation of sleep. Together with previous findings^5^, our results suggest that the multiplexed effects of different neuromodulatory systems, such as NE and ACh, may provide flexible regulation of local and long-range functional network interactions in the cortex that underlie state- and context-dependent behavior.

## Author Contributions

CB, FAC, and JAC designed the experiments. CB, FAC, and PX collected the data. CB analyzed the data. CB and JAC wrote the manuscript.

## Acknowledgements

The authors thank all members of the Higley and Cardin laboratories for helpful input throughout all stages of this study. We thank Rima Pant for generation of AAV vectors and assistance with immunohistochemistry and Dr. Andrew Moberly for assistance with the ex vivo imaging preparation. We thank the GENIE Project for jRCaMP1b plasmids and the lab of Dr. Yulong Li for the initial gift of Ne2m and rACh0.5 plasmids. This work was supported by funding from the NIH (R01EY022951, R01MH113852, R21EY036254, and R21AG087186 to JAC, F31 NS129354 to CB, and EY026878 to the Yale Vision Core) and a McKnight Scholar Award to JAC.

## Data Availability Statement

The full datasets generated and analyzed in this study are available from the corresponding authors on request.

## Code Availability Statement

Custom written MATLAB and Python scripts used in this study are available: https://github.com/cardin-higley-lab/Barnes_et_al_2026.

---

**Supplementary Figure 1.**
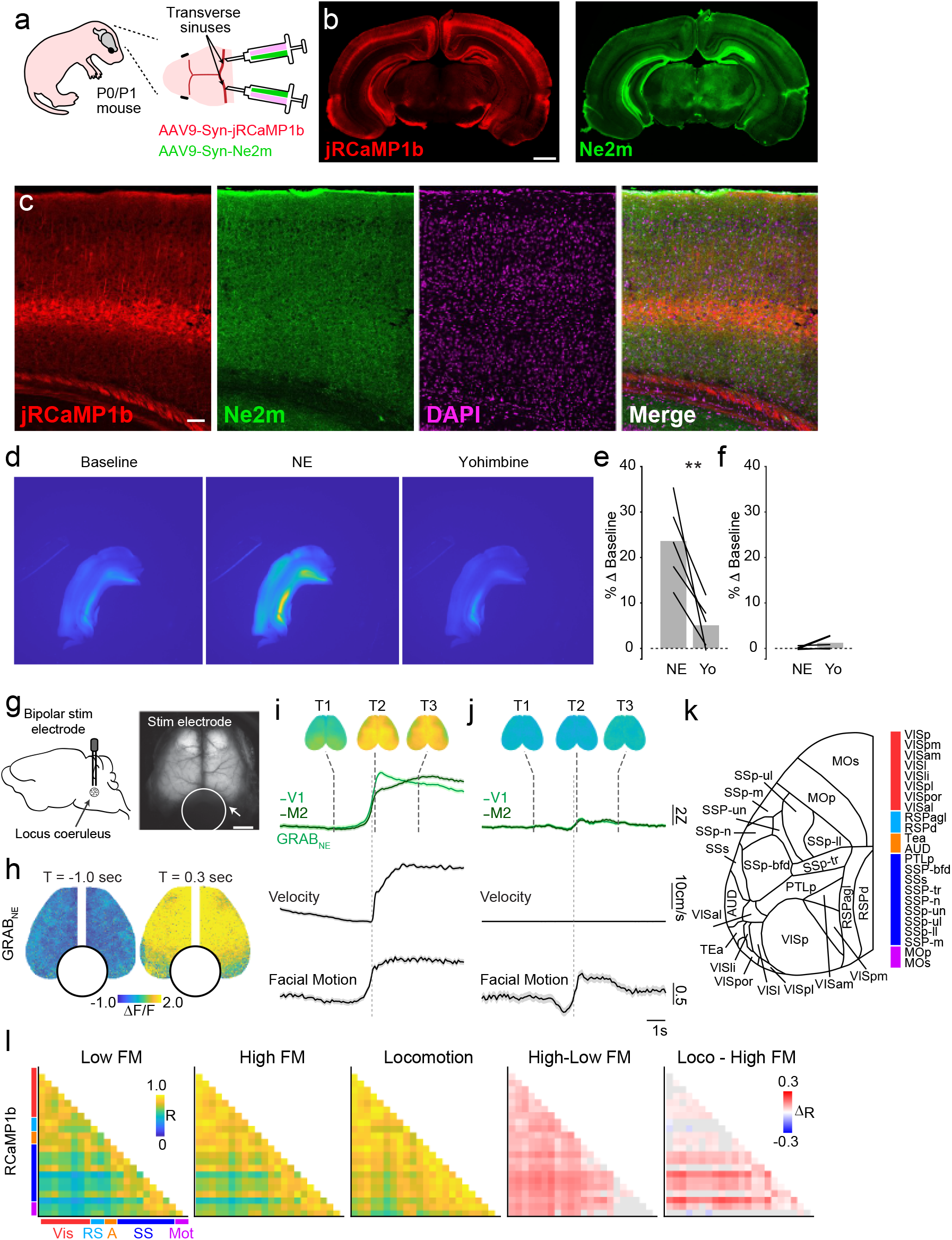
Validation of noradrenergic signaling and related controls. **a**, Schematic of the neonatal sinus injection approach. **b**, Example coronal fluorescent images from an adult mouse expressing jRCaMP1b (red) and GRAB_NE_ (green). Scale bar; 1mm. **c**, Example confocal images from an adult mouse co-expressing jRCaMP1b (red) and Ne2m (green) and stained for DAPI (magenta). Scale bar: 100 µm. **d**, Representative brain slice images from tissue expressing the GRAB_NE_ indicator during baseline conditions, during norepinephrine application (0.2 µM, equilibrium), and during subsequent yohimbine application (0.8 nM, equilibrium) conditions. **e**, Quantification of GRAB_NE_ signal as percent change from baseline within an ROI for norepinephrine (NE) and yohimbine (Yo) conditions under 470 nm illumination (paired t-test, n=5, P = 0.0039). **f**, Same analysis as in **e** but for 395 nm illumination (n=5, paired t-test, P = 0.2691). **g**, Schematic of bipolar stimulation electrode placement targeting the locus coeruleus and dorsal cortex view showing the electrode. **h**, Snapshots of GRAB_NE_ signals across the cortex prior to (T = -1s) and just after (T = 0.3s) electrical stimulation of the locus coeruleus. **i**, Event-triggered averages of jRCaMP1b activity from V1 and M2 cortex, aligned to locomotion onset. Top, snapshots of average responses at −2.5 s, +0.1 s, and +2.5 s. Middle, example V1 and M2 traces aligned to locomotion onset. Bottom, averaged wheel velocity and facial motion. **j**, Same as in **i**, but aligned to facial motion onset. **k**, Schematic illustrating Allen CCFv3 parcels, corresponding abbreviations, and color codes used in main and supplementary figures. **l**, Calcium correlation matrices for sustained low facial motion (Low FM), high facial motion (High FM), and locomotion states (n = 6 mice. Difference plots to the right show the change in correlations across states (stepwise significance assessed using stratified permutation testing (10K permutations) and Benjamin-Yekutieli FDR correction). Red indicates increased correlations, blue indicates decreased correlations, and gray indicates no significant change.

**Supplementary Figure 2.**
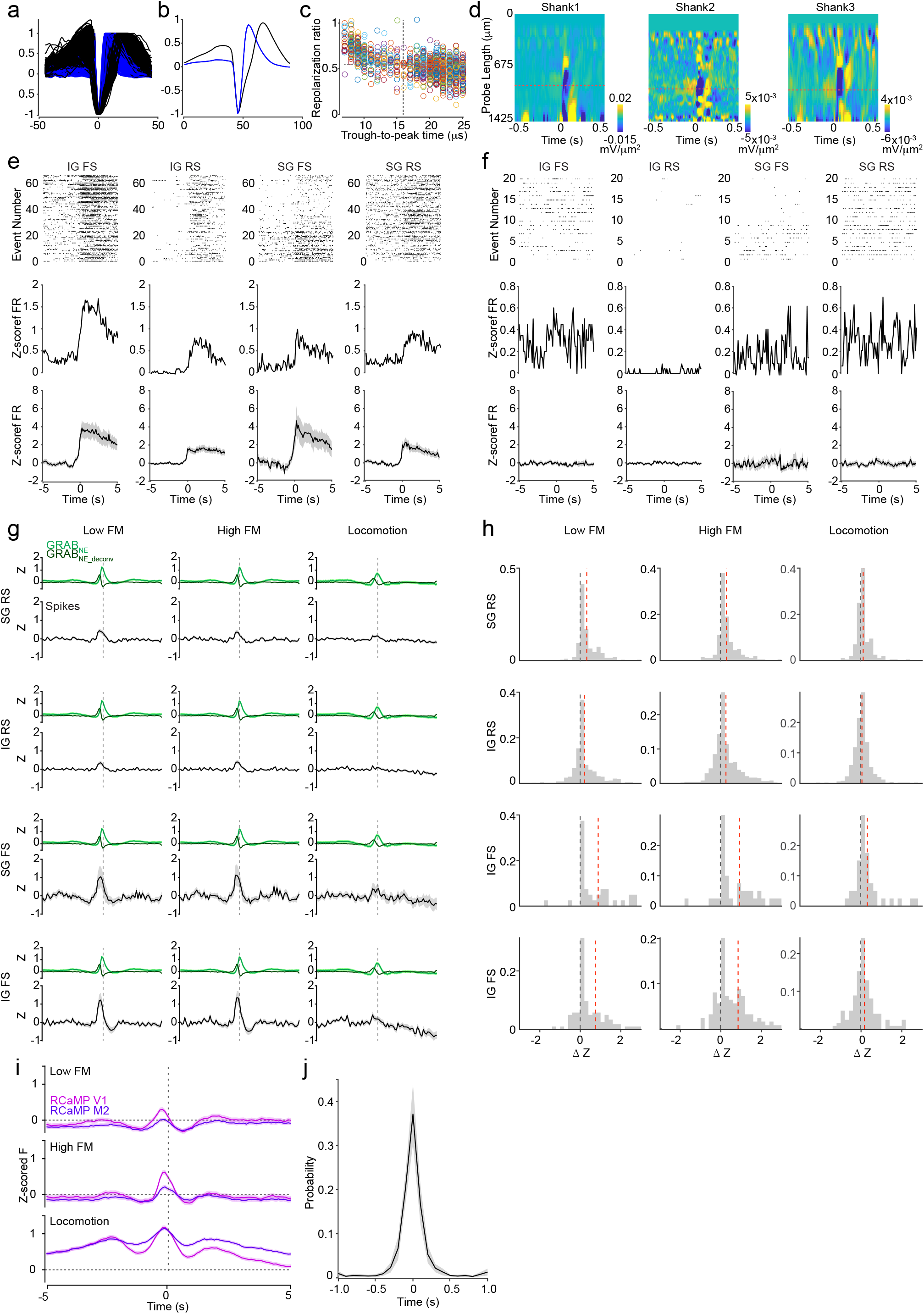
Classification and state-dependent modulation of cortical unit activity. **a**, Normalized spike waveforms for all recorded units. Waveforms were scaled to −1 at the trough and +1 at the post-trough peak. Regular-spiking (RS) units are shown in black and fast-spiking (FS) units in blue (n = 682 RS and 164 FS cells). **b**, Mean normalized waveforms for putative excitatory (RS) and inhibitory (FS) units. **c**, Scatter plot of repolarization ratio (measured at 1/3 ms) versus trough-to-peak time. Dashed lines indicate classification boundaries used to define cell types (FS: trough-to-peak ≤ 0.53 ms and repolarization ratio ≥ 0.55; RS otherwise). **d**, Example current source density (CSD) plots for three Neuropixel 2.0 shanks in one experiment. Layer 4 was identified by the prominent sink (blue) in each case. **e**, Locomotion-aligned activity by cell type. Columns correspond to infragranular (IG) FS, infragranular RS, supragranular (SG) FS, and supragranular RS units. Top row, example unit raster plots aligned to locomotion onset. Middle row, corresponding binned firing rates (125 ms bins) normalized using a regularized z-score (regularization factor = 0.1). Bottom row, population-averaged regularized z-scores. **f**, Same as in e, but aligned to facial motion onset. **g**, Population-averaged regularized z-scored firing for each cell type, aligned to NE peaks. Data are shown separately for low facial motion (Low FM), high facial motion (High FM), and locomotion states. Aligned NE fluorescence (GRAB_NE_) and deconvolved signals (GRAB_NE-deconv_) are shown alongside firing rate responses. **h**, Histograms of changes in regularized z-scores across cell types and behavioral states. No change is denoted by a black dotted line and the mean of the distribution is denoted by a red dotted line. **i**, Mean jRCaMP1b responses in V1 (magenta) and M2 (purple) aligned to GRAB_NE_ peaks in V1 during low facial motion, high facial motion, and locomotion states. **j**, Cross-correlogram of GRAB_NE_ peak timing between left and right V1.

**Supplementary Figure 3.**
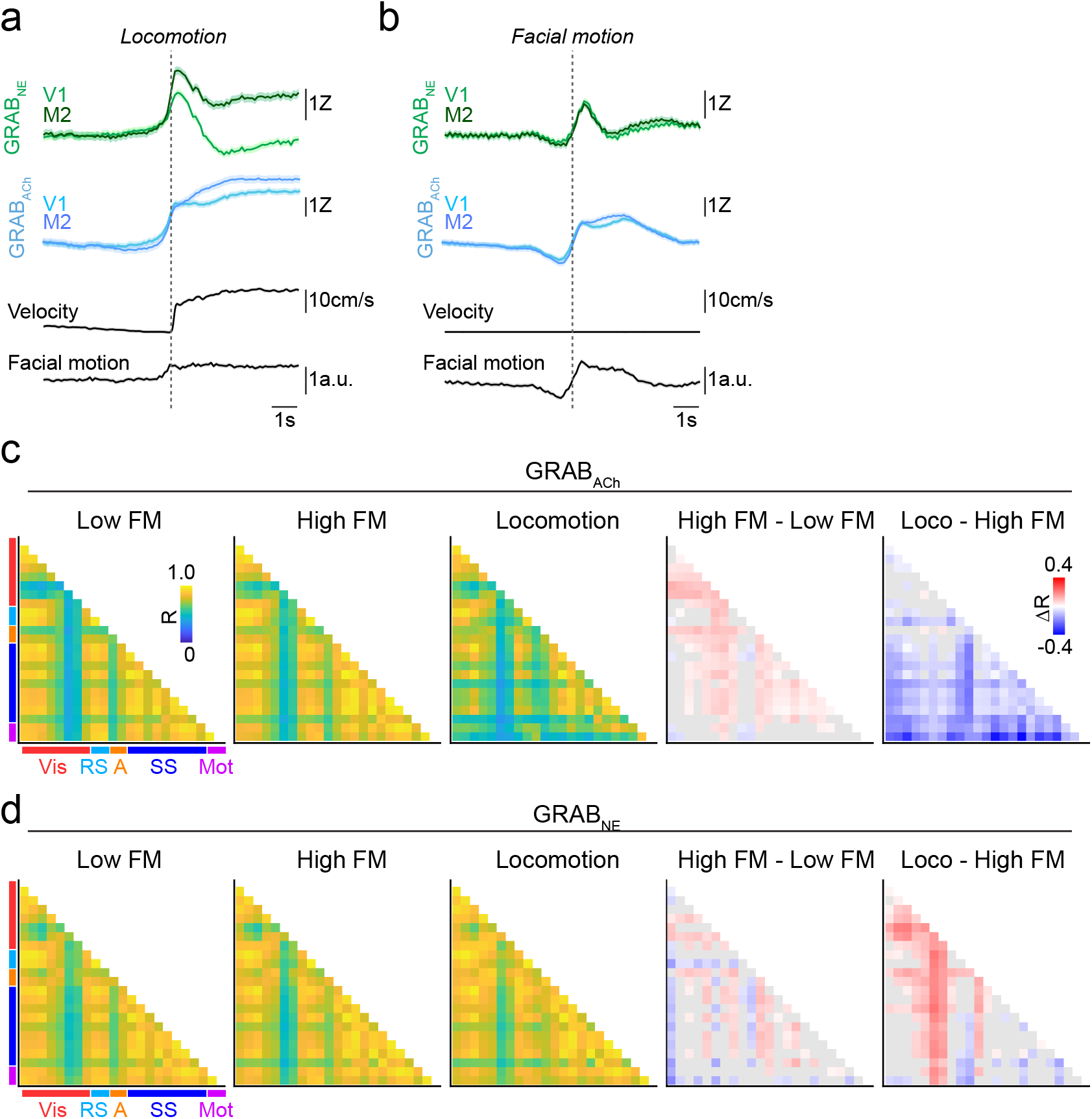
State-dependent dynamics and network structure of noradrenergic and cholinergic signals. **a**, Locomotion-aligned GRAB_NE_ (green) and GRAB_ACh_ (cyan) signals for norepinephrine and acetylcholine, respectively, in V1 and M2. Traces are aligned to locomotion onset (dotted line), with wheel velocity and facial motion shown below. **b**, Same as in a, but aligned to facial motion onset. **c**, Parcel-wise correlation matrices of GRAB_ACh_ activity during sustained low facial motion (Low FM), high facial motion (High FM), and locomotion states. Difference plots to the right show the change in correlations across states (n = 6 mice; stepwise comparisons (high facial motion vs low facial motion; locomotion vs high facial motion) assessed using stratified permutation testing (10K permutations) with FDR correction). Red indicates increased correlations, blue indicates decreased correlations, and gray indicates no significant change. **d**, Same as in **c**, but for GRAB_NE_ signals.

**Supplementary Figure 4.**
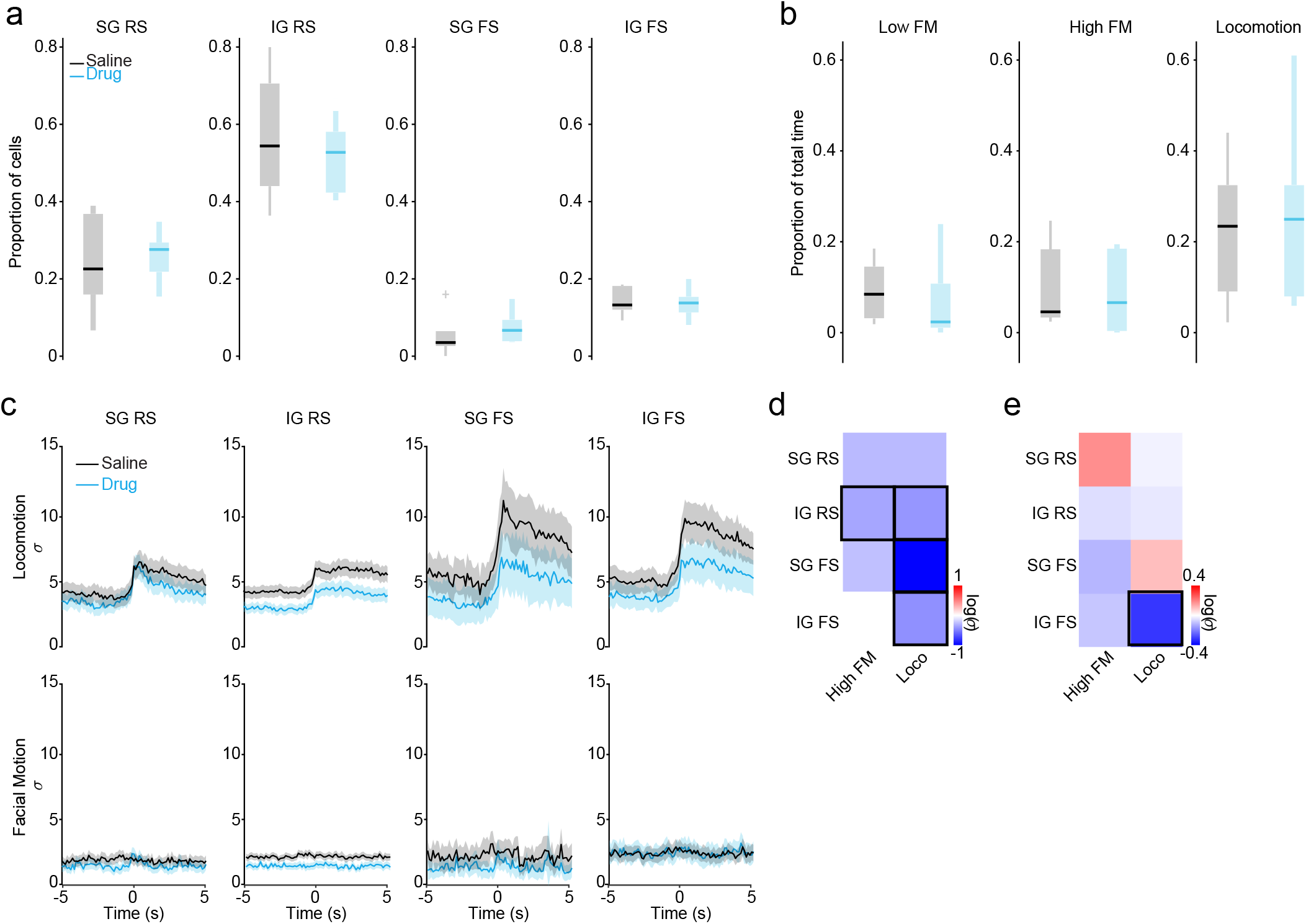
Control analyses for pharmacological manipulation experiments. **a**, Proportion of recorded units by cell type (supragranular (SG), infragranular (IG), regular-spiking (RS), and fast-spiking (FS)) under vehicle (black) and drug (blue) conditions. **b**, Fraction of time spent in low facial motion (Low FM), high facial motion (High FM), and locomotion states for vehicle and drug conditions. **c**, Average spiking activity of each cell type, aligned to locomotion (upper) and high facial motion (lower) onset under vehicle (black) and drug (blue) conditions. Data are normalized by the standard deviation of baseline activity during the sustained low facial motion state. **d**, Change in firing rate for each cell type during the baseline periods immediately preceding bouts of high facial motion (High FM) or locomotion (Loco) during drug application. Red indicates increased firing, blue indicates decreased firing, and black borders indicate significant changes. **e**, Change in the modulation of firing at the onset of facial motion (High FM) or locomotion (Loco) during drug application, calculated as an index value. Red indicates increased firing rate modulation, blue indicates decreased firing rate modulation, and black borders indicate significant changes.

**Supplementary Figure 5.**
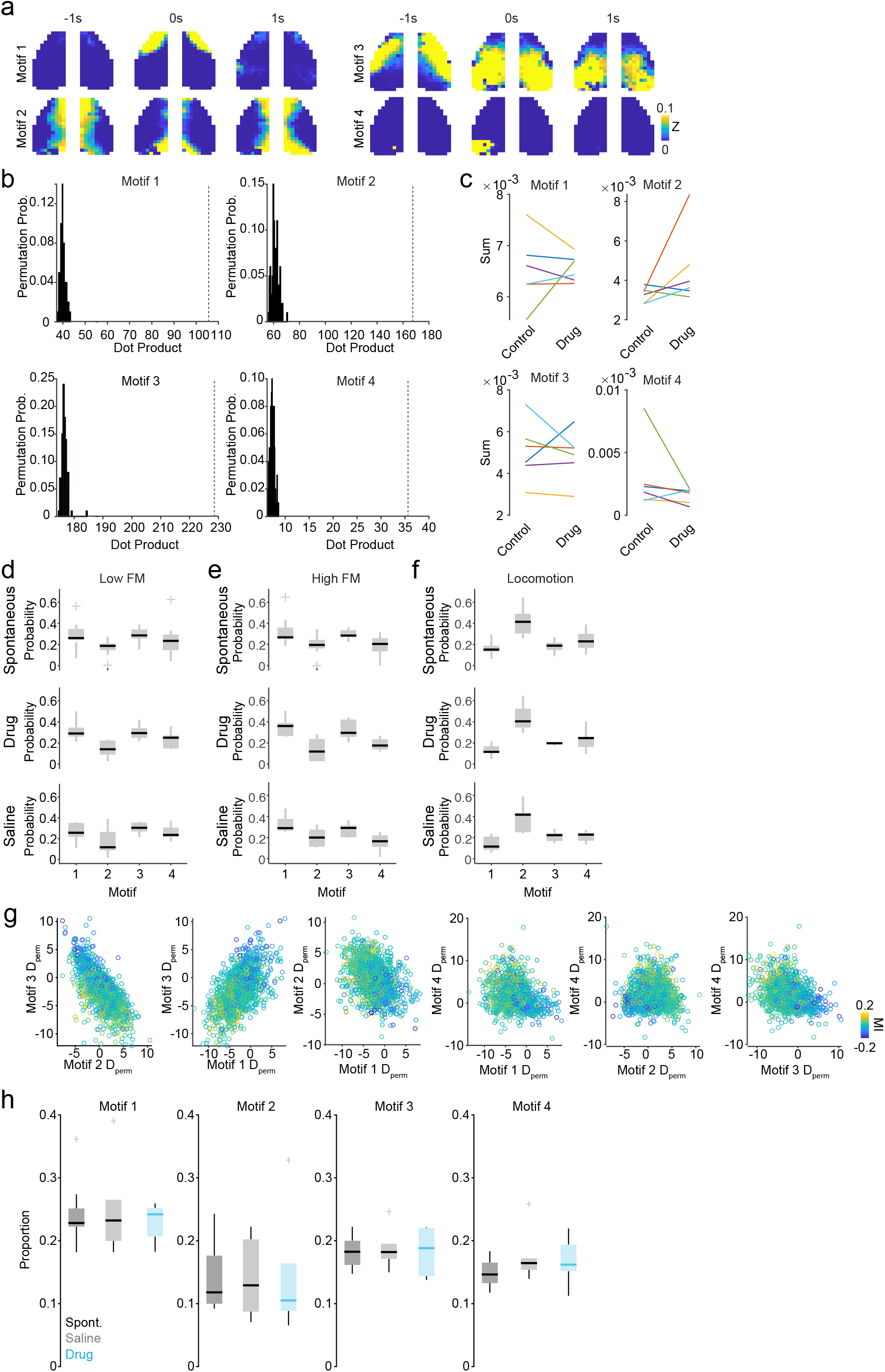
Large-scale motifs of cortical activity. **a**, Spatiotemporal progression of four common cortical motifs observed in all animals. For each motif, the activity pattern is shown at -1s, 0s, and 1s, with 0s representing the midpoint of the sequence. **b**, Motif validation using held-out data. Histograms show dot products between reconstructed activity from 10% held-out data and ground truth, compared against circularly permuted controls (100 iterations). Dashed vertical lines indicate true values. **c**, Mean magnitude of temporal motif weighting values per second for each motif, comparing control and drug conditions. **d**, Density of occurrence for motifs identified during the low facial motion state, shown as box-and-whisker plots across spontaneous, adrenergic receptor antagonist cocktail (drug), and vehicle conditions. **e**, Same as in d, but for the high facial motion state. **f**, Same as in e and e, but for the locomotion state. **g**, Tuning of individual units to motif combinations, quantified using a permuted Cohen’s D. Points are colored by each unit’s modulation index to norepinephrine peaks. **h**, Proportion of time spent in each motif during spontaneous activity periods (dark gray), vehicle (light gray), and drug (blue) conditions.

**Supplementary Figure 6.**
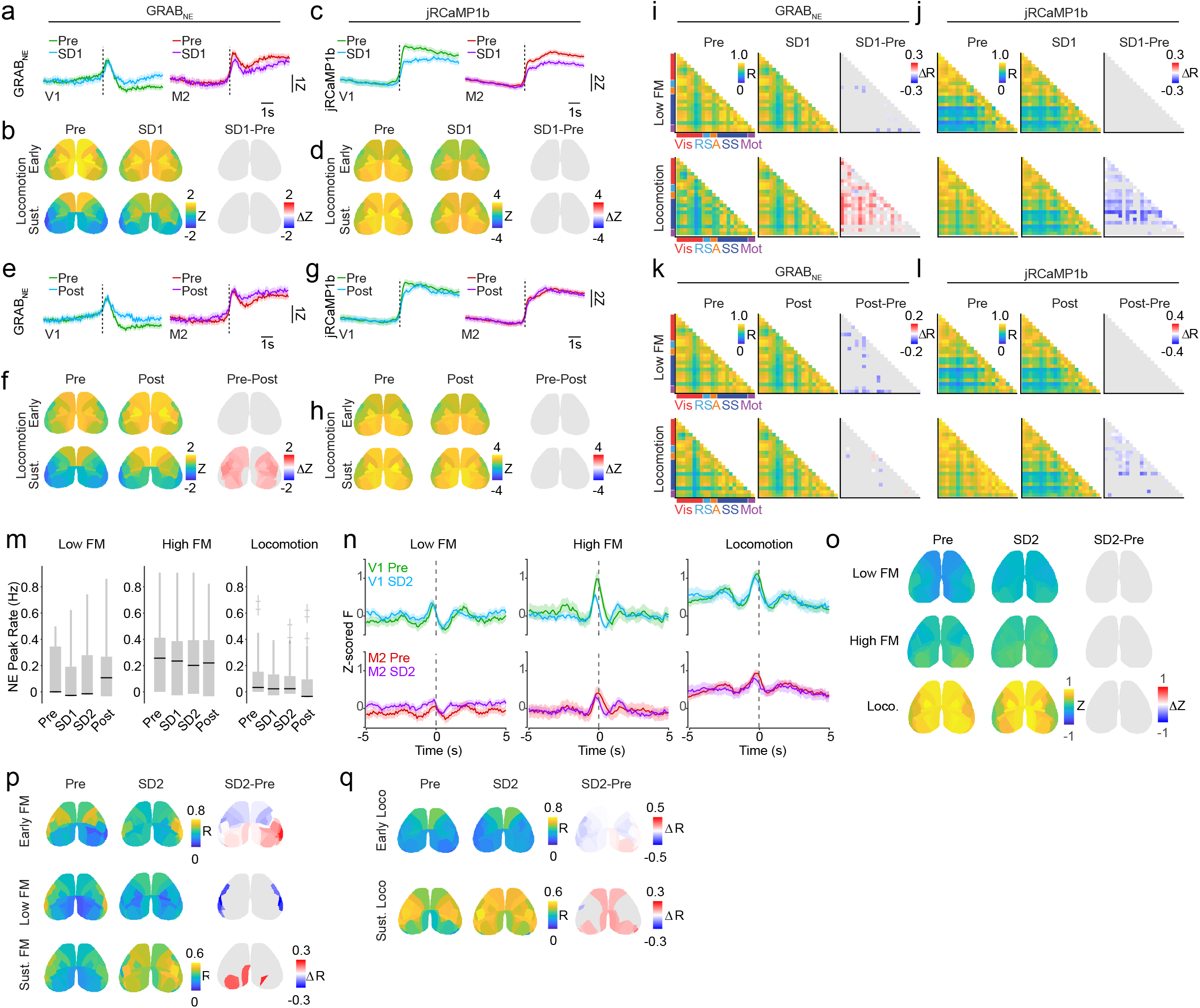
Sleep deprivation–associated changes in noradrenergic and neural dynamics. **a**, GRAB_NE_ signals from V1 and M2 aligned to locomotion onset (vertical dotted line), comparing sleep deprivation day 1 (SD1, cyan, purple) to the pre-deprivation baseline (Pre, green, magenta). **b**, Allen-parcellated dorsal cortex maps of GRAB_NE_ activity during early and sustained locomotion on days Pre and SD1. Difference map showing change in signal between the two days (SD1-Pre) is shown to the right. **c**, Same as in **a**, but for jRCaMP1b calcium signals. **d**, Same as in **b**, but for jRCaMP1b signals. **e**, GRAB_NE_ signals from V1 and M2 aligned to locomotion onset (dotted line), comparing recovery day (Post) to pre-deprivation baseline (Pre). **f**, Allen-parcellated GRAB_NE_ activity during early and sustained locomotion on days Pre and Post. **g**, Same as in **e**, but for jRCaMP1b signals. **h**, Same as in **f**, but for jRCaMP1b signals. **i**, Parcel-wise correlation matrices of GRAB_NE_ activity during low facial motion (Low FM) and locomotion, comparing sleep deprivation day 1 to baseline. Difference plot showing change across days (SD1-Pre) is shown to the right. **j**, Same as in **i**, but for jRCaMP1b signals. **k**, Parcel-wise GRAB_NE_ correlation matrices comparing post-deprivation recovery (Post) to baseline (Pre) for low facial motion and locomotion. Difference plot showing change across days (Post-Pre) is shown to the right. **l**, Same as in **k**, but for jRCaMP1b signals. **m**, Rate of GRAB_NE_ peak occurrences across sleep deprivation days and behavioral states. **n**, jRCaMP1b responses in V1 and M2 aligned to GRAB_NE_ peaks across behavioral states (Low FM, High FM, Locomotion) on days Pre (green, magenta) and SD2 (cyan, purple). **o**, Allen-parcellated jRCaMP1b activity during GRAB_NE_ peaks for sleep deprivation day 2 (SD2) and baseline (Pre), shown separately for each behavioral state. **p**, Parcel-wise correlations between GRAB_NE_ and jRCaMP1b signals during low facial motion (Low FM), early facial motion (Early FM), and sustained facial motion (Sust. FM), comparing sleep deprivation day 2 (SD2) to baseline (Pre). Difference plots showing the change across days are shown to the right (SD2-Pre). **q**, Parcel-wise correlations between GRAB_NE_ and jRCaMP1b during early (Early Loco) and sustained locomotion (Sust. Loco), comparing sleep deprivation day 2 (SD2) to baseline (Pre). Difference plots showing the change across days are shown to the right (SD2-Pre).

## Methods

### Animals

Male and female C57BL/6J, mice (2–5 months) were kept on a 12-h light/dark cycle, provided with food and water ad libitum and group housed when available following headpost implantations. Imaging experiments were performed during the light phase of the cycle. All animal handling and experiments were performed according to the ethical guidelines of the Institutional Animal Care and Use Committee of the Yale University School of Medicine.

### Neonatal sinus injections

Brain-wide expression of the NE sensor Ne2m and the calcium indicator jRCaMP1b or ACh sensor rACh0.5 was achieved by postnatal sinus injection as previously described ^100^. Specifically, P1 litters were removed from their home cage and placed on a heating pad. Pups were kept on ice for 3 min to induce anesthesia via hypothermia and then maintained on a metal plate surrounded by ice for the duration of the injection. Under a dissecting microscope, a small incision was made in the skin over the transverse sinuses. Viral injections were made with a NanoFil (WPI) attached to a 36-gauge needle and an UltraMicroPump (WPI) mounted to a stereotaxic arm. The needle was slowly lowered through the skull into the underlying transverse sinus. Pups were injected bilaterally with a 4 μL 50:50 mixture of AAV9-hSyn-Ne2m (1.8 × 1013 gc ml−1) and either AAV9-hsyn-NES-jRCaMP1b (2.5 × 1013 gc ml−1; Addgene) or AAV9-hsyn-rACh0.5 (1.8 × 1013 gc ml−1) per hemisphere. Viruses were injected at 1.2 μL/min, and the needle was left in the sinus for 30 s following the injection. Incision sites were sealed with Gluture glue, and pups were moved to a heating pad. Once the entire litter was injected, pups were gently rubbed with home cage bedding and nesting material and returned to their home cage.

### Surgical procedures

All surgical implantation procedures were performed on adult mice (>P50). Mice were anesthetized using 1–2% isoflurane and maintained at 37 °C for the duration of the surgery. The skin and fascia above the skull were removed from the nasal bone to the posterior of the intraparietal bone and laterally between the temporal muscles. The surface of the skull was thoroughly cleaned with saline, and the edges of the incision were secured to the skull with Vetbond. A custom titanium headpost was secured to the posterior of the nasal bone with transparent dental cement (Metabond, Parkell), and a thin layer of dental cement was applied to the entire dorsal surface of the skull. Next, a layer of cyanoacrylate (Maxi-Cure, Bob Smith Industries) was used to cover the skull and left to cure approximately 30 min at 22–24C room temperature to provide a smooth surface for transcranial imaging.

For simultaneous Neuropixel 2.0 and mesoscopic imaging experiments, a craniotomy was made over V1 as determined by a previously acquired retinotopic map. Quickcast was used to cover the hole before and between recording sessions. For simultaneous locus coeruleus stimulation in a subset of animals, stainless steel bipolar stimulating electrodes (125-μm diameter, Invivo1) were implanted at the following coordinates (anterior–posterior (AP) = −5.4 mm, mediolateral (ML) = -0.8 mm, dorsoventral (DV) = 4 mm, angle = 0 to target the locus coeruleus in the left hemisphere.

### Widefield imaging

Widefield calcium, noradrenergic, and cholinergic imaging was performed using a Zeiss Axiozoom with a PlanNeoFluar Z ×1, 0.25 numerical aperture objective with a 56-mm working distance. Interleaved epifluorescent excitation was provided by an LED bank (Spectra X Light Engine, Lumencor) using three output wavelengths: 395/25 nm, 470/24 nm and 575/25 nm. Emitted light passed through a dual-camera image splitter (TwinCam, Cairn Research) and then through either a 525/50-nm or 628/40-nm emission filter (Chroma) before it reached two scientific CMOS (sCMOS) cameras (Orca-Flash V3, Hamamatsu). Images were acquired at 512 × 512 resolution after 4× pixel binning, and each channel was acquired at 10 Hz with 20-ms or 35-ms exposure for dual and single channel recordings, respectively. Images were saved to a solid-state drive using HCImage software version 4.5.1.3 (Hamamatsu).

All imaging was performed during the second half of the light cycle in awake, behaving mice that were head-fixed so that they could freely run on a cylindrical wheel. An optical rotary encoder (Vex robotics) attached to the wheel continuously monitored wheel rotation. Mice received a multiday incremental handling and wheel-training habituation schedule (10 days: 3 days of handling, 1 minute on wheel with headpost grab desensitization without head fixation; and then 0.5-, 1-, 5-, 15-, 30-, and 60-min. head fixation) before imaging to ensure consistent running bouts. During widefield imaging sessions, the face (including the pupil and whiskers) was illuminated with an infrared LED bank and imaged with a miniature CMOS camera (Blackfly s-USB3, Flir) with a frame rate of 10 Hz. For mesoscopic imaging experiments acquired simultaneously with Neuropixel probe recordings, a synchronization pulse drawn from a lognormal distribution was sent to both acquisition systems for post hoc alignment.

### Retinotopic Mapping

Retinotopic mapping was performed by imaging jRCaMP1b at 20Hz with constant 575/25 nm LED illumination. Drifting horizontal or vertical bars of filtered noise at a variety of spatial and temporal frequences ^101^ were each displayed for 15 presentations. Mesoscopic data was averaged to the onset and duration (forced to be an odd number of samples) of horizontal and vertical stimuli presentations. The phase for the 1^st^ Fourier component was extracted for each pixel and stimulus orientation, to which a gradient was fit. The orientation of the gradient was found via the arctangent 2 function for each stimulus orientation. The resultant maps were combined by computing their difference and mapping to the complex plane. Sine maps were extracted by calculating the sine of the difference in angle encoded by each complex number.

### Electrophysiology

Electrophysiological data was collected with a Neuropixel 2.0 dense silicon probe ^102^, National Instruments chassis (PXIe-1071), and Open Ephys (v0.6.7) acquisition software. To minimize visual obstruction from the overhead mesoscopic imaging camera, the probe was inserted at a 45-degree angle with a Siskiyou micromanipulator (DR1000) at 5 μm/s. After 100 μm initial probe insertion, the reference was placed above the probe but continuous with the saline solution with a manual stereotaxic manipulator (Kopf). The burr hole was then covered with silicone oil (Thomas RECORDING GmBh, M-1000) to prevent evaporation and maintain conductivity. Activity was recorded along three shanks with the central shank densely sampled and flanking shanks configured into a tetrode recording pattern. Electrophysiological data was aligned to mesoscopic data through a microcontroller mediated synchronization pulse drawn from a lognormal distribution.

### Sleep Deprivation

Mice were sleep deprived for two consecutive days by preventing sleep during the light period. Mice were prevented from sleeping for the entirety of the light period (12 hours) on each day by periodically introducing novel objects (tubing, housing, food) and cage changes ^103-105^. Mice were monitored continuously for the 12-hour period. In the event that mice became unresponsive to novel object stimulation, gentle handling or stroking with a cotton-tipped swab was employed to prevent sleep. Mice were allowed to sleep ad lib during the following dark period.

### Histology

Histological validation was performed on a subset of animals at the conclusion of imaging experiments. Mice were deeply anesthetized with isoflurane and perfused transcardially with PBS followed by 4% paraformaldehyde in PBS. Brains were postfixed overnight at 4° and embedded in 1% agarose, and 50-μm sagittal sections were cut on a vibratome (VT1000, Leica. Slices were mounted on glass slides in Vectashield antifade mounting medium (Vector Laboratories). Widefield and confocal images were taken with a Zeiss LSM 900.

### Ex vivo imaging

Under isoflurane anesthesia, mice were decapitated and coronal slices (∼300 μm thick) were cut in ice-cold external solution containing (in mM) 100 choline chloride, 25 NaHCO3, 1.25 NaH2PO4, 2.5 KCl, 7 MgCl2, 0.5 CaCl2, 15 glucose, 11.6 sodium ascorbate and 3.1 sodium pyruvate, bubbled with 95% O2 and 5% CO2. Slices were transferred to artificial cerebrospinal fluid (ACSF) containing (in mM) 127 NaCl, 25 NaHCO3, 1.25 NaH2PO4, 2.5 KCl, 1 MgCl2, 2 CaCl2 and 15 glucose, bubbled with 95% O2 and 5% CO2. After an incubation period, slices were moved to a modified recording chamber under the objective of the widefield microscope and constantly perfused with oxygenated ACSF. Slices were imaged using the same protocol as during in vivo imaging sessions (10 Hz, 20-ms exposure). Norepinephrine and yohimbine were added to the perfusate.

### Pharmacology

All pharmacological experiments were performed with the experimenter blinded to the treatment and counterbalanced for the order of vehicle and drug administration. The craniotomy used for silicon probe insertion was filled with an adrenergic antagonist cocktail comprised of 0.5 mM each of yohimbine, propranolol and prazosin (Tocris Bioscience) or vehicle (0.1M PBS). The solution was allowed to diffuse for 45 minutes before data acquisition.

### Data analysis

All preprocessing and analyses were conducted using custom-written scripts in MATLAB (Mathworks, 2022b) or Python.

### Statistics and reproducibility

All statistical analyses were conducted using custom-written scripts in MATLAB and Python. All statistical results are listed in Supplementary Table 1. Sample sizes were not statistically predetermined but are similar to those reported in previous publications.

Each spontaneous behavior experiment was conducted in two sessions per animal, and analyses were performed on data averaged between sessions, except for stratified permutation tests for correlations where state data were pooled across sessions for each animal. Experiments were not independently replicated. The order of drug and vehicle sessions within each pharmacology group was counterbalanced. No animals were excluded from other analyses.

### Preprocessing of imaging data

Images with a size of 512 × 512 pixels were binned to 256 × 256 pixels, and frames were grouped by excitation wavelength (395 nm, 470 nm and 575 nm). For dual-color imaging, green and red images were acquired using different cameras and registered using a predetermined affine transformation. For each pixel, a 2^nd^ order Butterworth filter with a 0.001-Hz frequency cutoff was applied both forwards and backwards (matlab filtfilt function) to perform 0 phase high-pass-filtering. To improve signal to noise for the Ne2m-rACh0.5h cohort, the detrended data was smoothed with a box filter with a width of 5×5×5 through space and time. Hemodynamic artifacts were removed by a spatial-based nuisance signal removal method as previously reported ^100^.

For each timepoint t, let

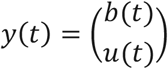

Where b(t) and u(t) are the demeaned target and nuisance signal, respectively, within a local patch of pixels. We assume the model:

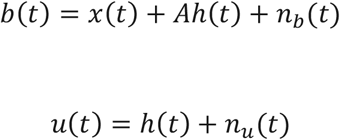

Where *x*(*t*) is the latent target dependent signal, *h*(*t*) is a shared nuisance signal, A is an unknown linear mixing operator, and *n*_*b*_(*t*), *n*_*u*_(*t*) are noise terms. The empirical covariance of the observations is

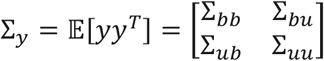

estimated from time samples. Measurement noise is modeled with variances 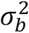 and 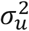, which in practice are determined by the median SVD. The covariance of the latent signal is estimated by the Schur complement:

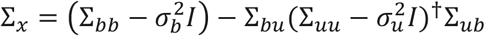

The latent target signal is recovered using the linear minimum least squared error (LMMSE) estimator

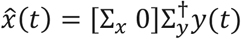

which yields the optimum linear estimator of *x*(*t*) given the joint target-nuisance observations under the assumed second order statistics.

For Ne2m-rACh0.5h cohorts, the noise estimate used in the hemodynamic correction model was spatially smoothed with 2D gaussian filter with a standard deviation of 2 to improve image quality. Images were registered to the CCFv3 using manually selected control points with instantaneous feedback on alignment quality via locomotion induced signal and retinotopic maps when available. For simultaneous Neuropixel and mesoscopic imaging experiments, vasculature was extracted from each session using the BCOSFIRE algorithm ^106^ and used to fit to the each subject’s retinotopic mapping session following an initial manual seed. The affine transformation matrices were then combined to enable a single transformation to the CCFv3 atlas. Time series for individual brain parcels were then extracted by averaging fluorescence values across all pixels within a parcel.

For experiments with simultaneous electrophysiology, data was aligned by taking the timing of the synchronization pulses detected by both acquisition systems, finding the peak cross correlation in their inter-event intervals, and fitting a linear model to transform timing data from the imaging system to the electrophysiology system’s sample numbers.

### Preprocessing of behavioral data

Wheel position was determined by the rotary encoder. To process optical rotary encoder data, the data was mapped to the 4 possible states of the sensor, then unwrapped and converted to centimeters given the radius of the wheel and rotational resolution of the sensor (2 degrees). The differential of the distance was smoothed by a quarter second gaussian window and converted to velocity given the sampling rate of data acquisition (5 kHz).

To identify bouts of locomotor activity, the velocity was converted to a binary signal via thresholding at 1 cm/s with rising and falling edges extracted as on and off timestamps, respectively. Locomotion bouts that had a peak velocity of less than 10 cm/s were removed. Bouts were merged if they occurred within 1 second of each other, and bouts that lasted for less than 5 seconds after merging were removed.

Facial motion was calculated by first extracting a mask of each subject’s face for each session with the segment anything tool ^107^. The convex hull of the logical mask was used to extract relevant pixels for subsequent processing. A SVD decomposition of the rectified and masked frame to frame change in luminance was computed, and the first principal component was taken as facial motion. Facial motion data was smoothed with a median filter window of .3 seconds, followed by moving average filter with a window of 1 second. Bouts of facial motion were extracted by thresholding the first principal component by the 60^th^ percentile determined during periods of no locomotion. Threshold cross events that occurred absent of locomotion that were at least 2s long were used for downstream analysis. Periods of low facial motion were similarly extracted from the bottom 40^th^ percentile.

### Preprocessing of electrophysiology data

Electrophysiology data was sorted with kilosort4 using the default parameters. To remove any impact of optoelectrical artifacts, detected spikes within 2 milliseconds of an LED state transition were discarded. The waveform for each cluster was computed by sampling up to 500 spikes per cluster, high passing with a (0 phase,3^rd^ order, 300Hz cut off) Butterworth filter, and calculating the mean electric potential across all channels centered in time around each spike within a 3 ms window. The waveform from the channel with the largest deflection was used to assess unit quality and cell type for each cluster. The false positive contamination rate was estimated by the number of spikes falling within the refractory period. The false negative contamination rate was determined by the amplitude cutoff in spikes. Clusters that did not surpass 10% false positive contamination, 10% false negative contamination, and whose waveform did not cross through excludable regions of time-voltage space were used in subsequent analyses. Units were classified as putative excitatory (regular spiking/pyramidal) and inhibitory (fast spiking/PV) based on the peak-trough and repolarization time of representative waveforms for each cluster ^3^. Briefly, the average normalized waveforms of all units were clustered with the k-means method based on 2 parameters: peak to trough time, and repolarization (i.e. defined as the value of the normalized waveform 0.45ms after peak). FS units had higher repolarization values and shorter peak-to-trough times than RS units.

The local field potential was averaged to full-field drifting grating visual stimuli presented before the onset of mesoscopic imaging. Current source density analysis was performed by taking the negative of the second derivative vertically across each shank. Layer 4 was manually identified as the location of earliest sink per shank. Clusters with a primary channel below or above the identified sink were classified as infragranular or supragranular, respectively ^108, 109^.

### Postprocessing analysis

Imaging data from individual parcels were used for subsequent analyses. The medial Allen Mouse Brain CCFv3 prelimbic (PL) and anterior cingulate area (ACA) parcels were removed due to their small size and prominent vasculature along the midline. Colliculus regions (superior colliculus (SC) and inferior colliculus (IC) were also removed, resulting in a final total of 23 Allen Mouse Brain CCFv3 parcels per hemisphere. The primary visual cortex (V1 or VlSp) and the secondary motor area in the frontal cortex (M2 or Mos) were selected as representative areas for some population summaries.

### Motif generation and analysis

To extract neural activity motifs, frames were median binned (8×8) to a final size of 32×32. To facilitate identifying common motifs across subjects and sessions, each pixel was Blom normalized. A softmax normalization was applied to each frame of the resultant data with a temperature parameter of 0.3. The normalized data was appended across subjects and motifs were extracted with the SeqNMF algorithm (K=15, L=31, *λ*=0.0005) ^70^. A custom regularization factor was implemented to encourage spatial spread in motifs, defined as follows. Let X be the data matrix SeqNMF factorizes as

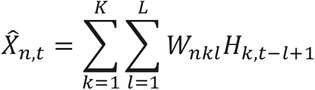

We define *W*_*nk*_ as

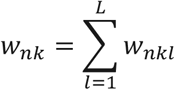

and the penalty term, *P*_*nk*_, as

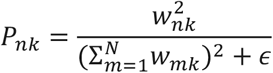

Which is included in the denominator of the update rule and scaled by ***λ***_*SpreadW*_, which was set to 1.

Motifs were validated by witholding 10% of each session and fitting the temporal loading while keeping the learned motifs fixed. The similarity of the reconstructed data to ground truth was assessed against reconstructions with circularly permuted temporal loadings. Finally, the temporal loadings for the whole dataset was fit and used for subsequent analysis.

The relationship between single cells and motifs was quantified by taking the cross-correlation of the temporal loading and binned spike rate for each motif, both normalized to to have a sum of 1. The -0.5s to +1.5s window of the cross-correlation was normalized by subtracting the median cross correlation value in the -2s to +2s window. A null distribution with 1,000 samples was established by circularly permuting the relationship between the binned spike rate and motif temporal loading. The mean and standard deviation of the null distribution was used to establish a Cohen’s D metric, which was taken as a measure of network tuning between each cell and motif. Similarly, we quantified the degree of modulation by NE for each cell by finding the number of spikes 1s before and after each peak in the NE signal and calculating Cohen’s D in the standard way. The motif and NE peak derived statistics were related through a linear mixed effects model of the form

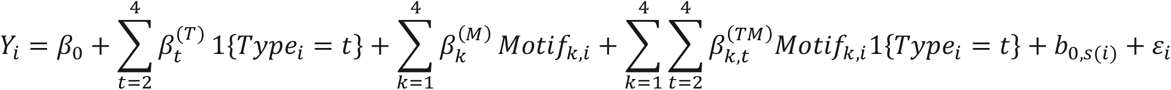

Where *Y*_*i*_ is NE peak modulation, Type is the cell type categorical variable (with *t* = 1 as the reference), and *Motif*_*k*_ is the motif tuning metric for observation *i* of subject *s*(*i*). The random effects are defined as

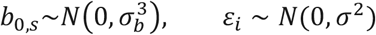

We extracted the fixed-effect coefficient vector 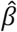 and its covariance matrix 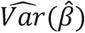 for each motif *k* ∈ {1, …,4} and each cell type *j* ∈ {1, …,4} we computed the Type-specific motif slope as a linear contrast:

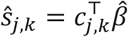

Where *c*_*j,k*_ is a contrast vector that selects the motif main-effect coefficient for *Motif*_*k*_, and for non-reference types, also selects the corresponding interaction coefficient *Type*_*j*_: *Motif*_*k*_, which yields

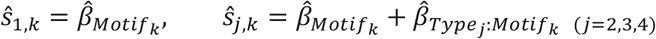

To assess if slopes differed from 0, we performed Wald tests using the matlab function “coefTest” followed by Benjamini–Yekutieli correction over all tests.

To examine the impact of pharmacological intervention on the relationship between motif tuning and NE modulation, we similarly fit a linear mixed effects model that included the condition as a categorical variable, allowing condition, type, and motif interactions:

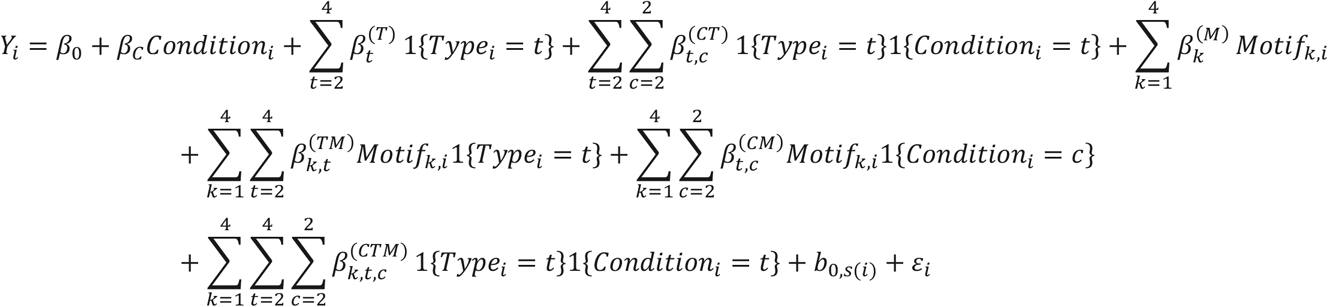

To assess if there was any effect of the condition, we fit a reduced model to compare:

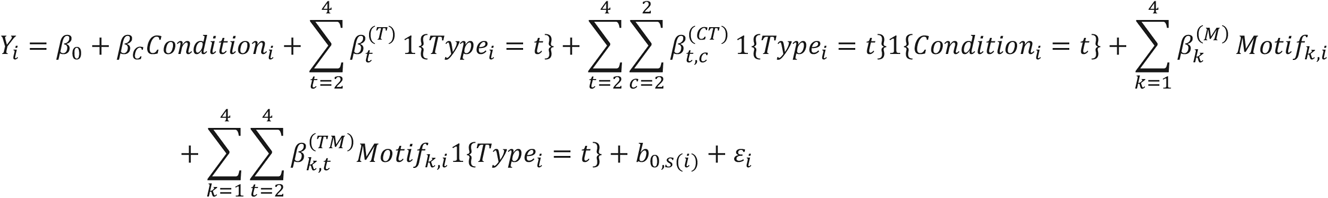

and tested the joint null hypothesis

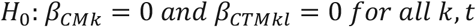

With the likelihood ratio test, which has a test statistic of

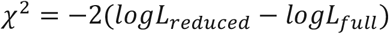

Where *L*_*reduced*_ and *L*_*full*_ are the likelihoods for their respective models. As the test statistic is asymptotically chi-square distributed, a significance level can be determined.

### Deconvolution

To remove the delay introduced by sensor kinetics, we deconvolved the Ne2m signal by its published time constants. We assume the observed signal is a linear convolution with a known impulse response function:

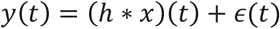

Where *y*(*t*) is the observed signal, *x*(*t*) is the latent signal, *h*(*t*) is the sensor convolution, * denotes the convolution operator, and *ϵ*(*t*) is measurement noise. The latent signal in the frequency domain was calculated as:

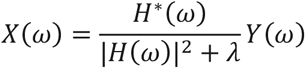

Where *H*^*^(*ω*) is the complex conjugate of *H*(*ω*) and *λ* is a small regularization constant to minimize the impact of noise (Tikhonov regularization). The time domain signal was obtained by taking the inverse fast Fourier transform of *X*(*ω*).

### Locus coeruleus stimulation-evoked activity

Locus coeruleus stimulation comprised a brief burst at 100 Hz (1-ms pulse width, 20 pulses, 60–100 μA). Trials in which stimulation caused locomotion were automatically excluded. Data were normalized by subtracting the baseline mean in the prestimulus period (−2 s to 0 s). Difference ΔF/F data were averaged across all trials within an animal and summarized as mean ± standard error of mean (s.e.m.) across animals.

### State transitions and sustained states

Behavioral states were categorized into high and low states (high or low facial movement and quiescence or locomotion) based on movement, as described above. Changes at transition from the low to high movement state (facial movement onset or locomotion onset) were defined as state transients. For locomotion, only trials that contain at least 5 s of running and, 2s or preceding were preceded by a minimum 2 s of quiescence, and reached a maximum velocity of 10 cm/s were included. For high facial movement state onset, if a high facial movement state was preceded by at least 4 s of a low facial movement state, the transition between the two states was used as the high state onset time point. To quantify state transition changes, the fluorescent values of each parcel were normalized (z-scored) by the pre-state period (-5 to -2 seconds before onset) for each bout of the state transition. The early and sustained changes were calculated as the mean z-scored values between 0 to 1 seconds after onset, and 2 to 3 seconds after onset, respectively. To test for significance, stratified permutation testing was performed with 10K permutations and Benjamini–Yekutieli false discovery rate correction. Raw and adjusted P values are available in Supplementary table 1.

### State-dependent correlations

Sustained state correlations between every pair of Allen Mouse Brain CCFv3 parcels in Ne2m, jRCaMP1b, or rACh0.5 signals were calculated with a minimum duration of 2 s . When comparing across days, the number of state epochs were matched for locomotion and high and low facial movement. Inter-signal correlations were calculated in a similar manner except that the correlations were performed between the two signals within a parcel. Correlation matrices were compared between locomotion and high facial movement states as well as between high facial movement and low facial movement states. For interventions across days, the same state was compared across experimental conditions. To assess the significance of differences in pairwise correlations, a stratified permutation test was used. Labels were shuffled within subject, and the mean correlation taken subject-wise. The difference in the mean correlation was generated 10,001 times to build a null distribution that was compared against the observed test statistic. Multiple-comparisons correction was performed with the fdr_bh.m toolbox in MATLAB with a false discovery rate (FDR) of q < 0.05 and the the Benjamini–Yekutieli method.

For the state transition correlation analysis, the time period -3s to 3s relative to the transition time was extracted, averaged per subject, and then z-scored. A linear mixed effects model was then fit to each parcel with the form

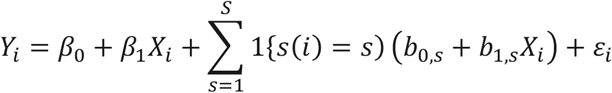

where *X*_*i*_ and *Y*_*i*_ are the two signals and *s* indicates the subject. A Wald test on the *β*_1_ coefficient provided a P value which was corrected for multiple comparisons with Benjamini–Yekutieli (q<0.05). All raw and corrected values are available in Supplementary table 1.

